# Mapping functional traces of opioid memories in the rat brain

**DOI:** 10.1101/2023.08.07.552221

**Authors:** Joana Gomes-Ribeiro, João Martins, José Sereno, Samuel Deslauriers-Gauthier, Teresa Summavielle, Joana E. Coelho, Miguel Remondes, Miguel Castelo Branco, Luísa V. Lopes

## Abstract

Addiction to psychoactive substances is a maladaptive learned behavior. Contexts surrounding drug use integrate this aberrant mnemonic process and hold strong relapse triggering ability. Here we asked where context and salience might be concurrently represented in the brain, and found circuitry hubs specific to morphine-related contextual information storage.

Starting with a classical rodent morphine-conditioned place preference (CPP) apparatus, we developed a CPP protocol that allows stimuli presentation inside a magnetic resonance imaging scanner, to allow the investigation of whole brain activity during retrieval of drug-context paired associations, as well as resting state functional connectivity under the effect of morphine conditioning. Using fMRI we found context-specific responses to stimulus onset in multiple brain regions, namely limbic, sensory, and striatal. Furthermore, we found increased functional connectivity of lateral septum with regions within and beyond a proposed limbic network, and of the lateral habenula with hippocampal CA1 region, in response to repeated pairings of drug and context.

Subsequent exposure to either morphine or saline-conditioned contexts led to significant, context-specific, functional interconnectivity among amygdala, lateral habenula, and lateral septum. Resting-state connectivity of the lateral habenula and amygdala, and that during saline-paired context presentation significantly predicted inter-individual CPP score differences.

In sum, our findings show that drug- and saline-paired contexts form distinct memory traces in overlapping functional brain microcircuits, and intrinsic connectivity of habenula, septum, and amygdala likely influences the maladaptive contextual learning in response to opioids. We identify functional mechanisms involved in the acquisition and retrieval of drug-related memories that might be behind the relapse-triggering ability of opioid-associated sensory/contextual cues.

**Graphical abstract:** To investigate the brain-wide regional activity underlying drug addiction Gomes-Ribeiro et al. have developed a rodent morphine-induced CPP (conditioned place preference) protocol whose contextual cues can be consistently presented inside an MRI apparatus. The authors found a common circuitry supporting the neural mechanisms responsible for memorizing an association between distinct contexts, and morphine or the absence thereof, that includes regions involved in affect, reward, contextual perception and memory, with subtle, intriguing, functional specificities ultimately underlying the storage of distinct, individual, CPP memory engrams across the different animals. Animals that are more prone to strong emotional responses (as measured by the baseline resting state amygdala and habenula functional connectivity), exhibit a kind of neural circuit priming effect, and thus develop stronger connectivity patterns in anticipation of stronger morphine addiction behavior. These findings could help to clarify the inter-individual sensitivity to opioids in humans, since despite responding positively to the effects of opioids, many humans do not develop an addiction.

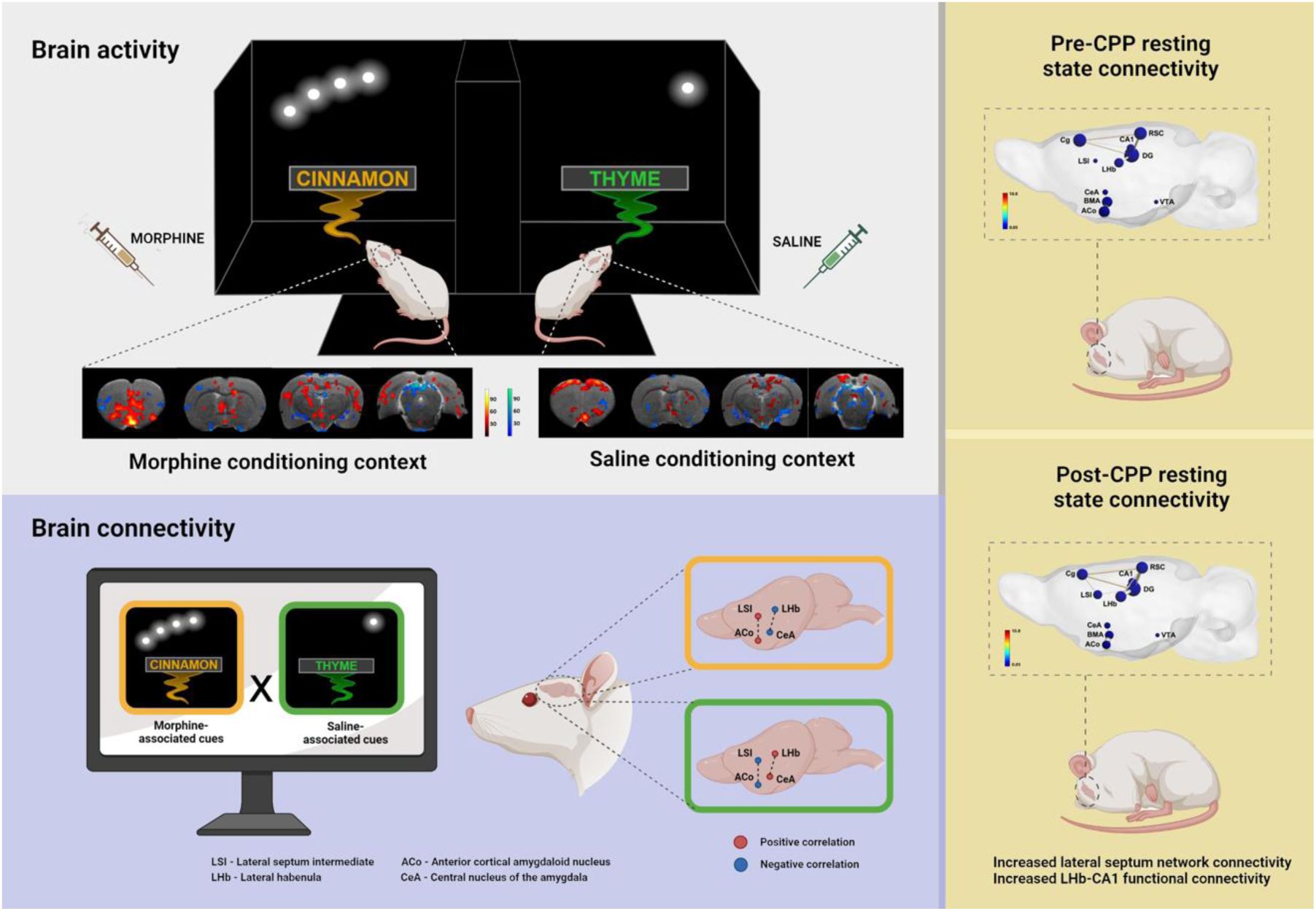

## INTRODUCTION

An ongoing opioid addiction epidemic currently threatens populations worldwide. Opioids are classical central nervous system depressants, and the most effective anti-nociceptive drugs, and create strong physical and emotional dependence, frequently leading to overdose (Hayes et al., 2020). Opioids bind to opioid receptors in local and efferent GABAergic neurons of the ventral tegmental area (VTA), leading to their hyperpolarization and silencing, in turn allowing dopaminergic (DA) neurons to switch from tonic to phasic firing modes (Bourdy & Barrot, 2012). Such change leads to a surge of dopamine release onto target regions of the mesocorticolimbic system (Barrot, 2015), causing an intense sense of euphoria, pleasure, and relaxation (Becerra et al., 2006; Nummenmaa & Tuominen, 2018). Such positive affect constitutes a strongly salient stimulus, the lack thereof produces severe withdrawal symptoms (e.g., dysphoria, pain, anxiety, and depression), both motivating and perpetuating drug seeking (Spanagel et al., 1992). These aversive physical and emotional states, stronger than those found with other types of drugs (e.g., amphetamine and cocaine) (Kutlu & Gould, 2016), can only be avoided, or relieved, by retaking the drug, further reinforcing the motivational value of opioids (Kenny et al., 2006; Pergolizzi et al., 2020). Thus, drug addiction is a maladaptive learning process.

Drug use becomes strongly associated with surrounding context, and emotional states become conditioned to the perception of environmental stimuli (Kenny et al., 2006; Pantazis et al., 2021). Contextual cues such as substance use paraphernalia, visual scenery, odors, and sounds, are encoded in the brain (Le Merrer et al., 2012) and become motivationally charged in strong associative memories (Fatseas et al., 2011). Once present, such stimuli can lead to the reactivation of these stable memories, and on their own drive opioid (re)use, even after extinction (Hearing et al., 2016). Thus, context plays a critical role in drug repeated use and relapse. In spite of intense research, the neural mechanisms responsible for maintaining and retrieving these maladaptive memories remain unclear.

The conditioned place preference (CPP) paradigm is commonly used to model drug-context associations. Here, the positive affect resulting from opioid-induced dopamine release becomes conditioned to the sensory (visual, odorant, and tactile) cues present in a given enclosed environment (Bardo & Bevins, 2000). Thus far, several interconnected regions of the mesocorticolimbic circuit have been implicated in the encoding and retrieval of opioid-induced CPP memories, including the VTA (Harris et al., 2004), nucleus accumbens (NAc) (Ma et al., 2009), prefrontal cortex (PFC) (Wang et al., 2019), amygdala (Amy) (Zarrindast et al., 2003) and the hippocampus (HIPP) (Kutlu & Gould, 2016). In natural environments, contextual information is initially represented in the hippocampus before being transferred to higher order cortical centers for long term memory storage (Preston & Eichenbaum, 2013). The strong reciprocal hippocampal-cortical connections, along with its numerous subcortical inputs and outputs, place the hippocampus as a hub for encoding and retrieval of drug-context memories, something that has been shown in early hippocampal lesion studies, with hipocampectomized rats unable of acquiring drug-induced place preference (Taubenfeld et al, 2010). Consistent with this, morphine-CPP retrieval was associated with increased basal synaptic transmission and impaired long-term potentiation in the HIPP (Portugal et al., 2014), and HIPP dentate gyrus (DG) granule cells, involved in context discrimination (Cholvin & Bartos, 2022), increase the expression of activity-related c-Fos on exposure to a morphine-related context (Rivera et al., 2015). Finally, reward expectancy in the morphine-CPP paradigm increased high frequency gamma oscillations in the ventral HIPP-NAc circuit (Sakae & Martin, 2019).

Despite numerous reports highlighting the representation of drug-associated contexts in the HIPP, recent contextual fear conditioning studies show that memory engrams are formed, and consolidated, in the HIPP and other connected structures such as mPFC and basolateral amygdala (BLA) (Kitamura et al., 2017). Such remote memory encoding is thought to support the stabilization of hippocampal representations of contexts before they gradually mature in the PFC (Marks et al., 2022). Given the extensive midbrain dopaminergic innervation of the HIPP, and its role in stabilizing neural maps biased towards particularly rewarding events (Gomperts et al., 2015; McNamara et al., 2014), VTA is another region likely playing a role in the regulation of encoding and reactivation of “drug maps”.

To identify the neural circuits underlying drug-induced memory storage and retrieval, we used a classical morphine-CPP apparatus, redesigned the conditioning contexts, and developed a protocol to present the conditioned context stimuli inside a rodent magnetic resonance imaging apparatus (MRI). For this we first paired two distinct contexts with an administration of either morphine (reward) or saline (neutral), thus attributing to the two contexts distinct affective salience. We then asked what brain regions concurrently represent contexts and saliences, under the assumption that although both contexts may be encoded by a single population of neurons, the overlay of a motivational value might engage specific neural assemblies within that population, each representing rewarded (morphine) and neutral (saline) contexts. Despite responding positively to the effects of opioids, many humans do not develop an addiction (Koob & Volkow, 2016). Seminal work with rodent models of morphine-CPP (Bardo & Bevins, 2000) also revealed contrasting subject-dependent levels of CPP expression. Our general prediction was that CPP expression levels would be linked to the strength of functional connectivity in memory-related and/or valence assignment regions during presentation of morphine-paired contexts, compared to the level of connectivity during exposure to saline-paired contexts. For rats that would not develop CPP, either the reverse or the absence of a correlational effect would be expected.

Once transferred to the MRI setup, the dominant contextual cues, olfactory and visual, proved sensitive enough to reflect inter-individual variation, with learner and non-learner animals. Thus-analyzed brain activity showed significant context-specific responses from limbic, sensory, and action initiation regions, co-occurring with the onset of cue presentation. Our data supports our prediction that functional connectivity in memory-related and/or valence assignment regions is linked to CPP expression level, but not during morphine-associated context retrieval. Lower functional connectivity during saline conditioning may lead to animals being less able to form distinct representations of the conditioning contexts and the emotional states therein. By analyzing neural co-activity in identified brain circuits supporting drug-related motivation during drug-reward conditioning, we provide insights into the long-term plasticity underlying drug use.

## METHODS

### Subjects

Animal procedures were performed at the Rodent Facility of Instituto de Medicina Molecular (iMM), licensed under the reference number 017918/2021, and at Institute for Nuclear Sciences Applied to Health (ICNAS) Pre-Clinical Facility, in compliance with the European Directive 2010/63/EU, transposed to Portuguese legislation in DL 133/2013. All animal research projects were reviewed by the Animal Welfare Bodies (ORBEA) of each institution and approved by national authority, to ensure animal use was in accordance with the applicable legislation and following the 3R’s principle. Male Sprague-Dawley rats (Charles River Laboratories, France) aged 12-16 weeks were used for all behavioral experiments (total n = 30). Environmental conditions were kept constant: food and water ad libitum, 22-24°C, 45-65% relative humidity, 12h light/12 dark cycles, and housed in groups of five, except for the MRI group which was housed in pairs.

### Drugs

Rats were administered either a morphine hydrochloride (5 mg/kg SC) or saline (0.9%) solution. For fMRI experiments, an induction dose of isoflurane (2-3%) and both bolus (0.05 mg/kg) and maintenance (0.1 mg/kg/hr) doses of Medetomidine were administered for light sedation. This is proven to be efficient in immobilizing the animal inside the MRI scanner for several hours, all the while maintaining neural activity and neurovascular coupling intact (Sirmpilatze et al., 2019). All procedures were performed in accordance with EU and Institutional guidelines.

### Behavioral experiments

#### Conditioned Place Preference adaptation

We aimed to develop a conditioning paradigm that could be effectively transferred to an MRI scanner. Selection of conditioning cues started with an assessment of spontaneous odor preference for a variety of odorants, as described in Lee et al. (2013) (Lee et al., 2013). Male Sprague Dawley rats (n=8) were isolated in a common rat cage and presented with herb and/or spice water infusions. A few mL was applied onto filter paper in an inverted small petri dish placed on top of the cage grid. We then timed active investigation, defined as periods in which the animal’s nose was within 1 cm of the petri dish, during a 120-s time lapse. Rats completed 6 trials in the 1^st^ day, and 7 on the 2^nd^ day. Odorant (table) presentation sequence was randomized across rats at 2-min intervals. We then normalized investigation times by dividing the time spent in the vicinity of an odorant, and the summed investigation times of all odorants on that day. This number was then divided by the water exploration time to get a normalized preference metric. Normalized preference was then compared across odorants using ANOVA followed by *post-hoc* comparisons (Fisher LSD). This study showed that basil and bay were the least “preferred” odorants. All other odorants were considered “neutral” for the purpose of this research (Supplementary figure 1-a).

The two odorants eliciting similar investigation times (cinnamon and thyme) were then transferred to the Conditioned Place Preference (CPP) behavioral apparatus. This consisted of a black Plexiglas box divided into three compartments: two main squared chambers of 40 x 40 x 40 cm, each opened by a 15 x 40 cm entrance, each containing one or the other odor cues, connected by an external neutral compartment of 40 x 15 x 40 cm. The animals were placed in the neutral compartment of the CPP apparatus, and the exploration times of the conditioning chambers were assessed for 15 minutes. Rats showed no preference for either context as exploration times were similar (Supplementary figure 1-c). Next, we coupled each of two presented odorants with visual cues composed of either one or a set of four aligned equidistant white LEDs at minimal intensity, which were placed above eye level on three out of the four black walls of the conditioning chambers. The rat’s visual system mainly allows the detection of light contrasts and intensities, and less individual colors (Jacobs et al., 2001). The single and 4-set LED visual stimuli were paired, respectively, with thyme- and cinnamon-odorized black foam sheets located near the base of the compartment walls. No additional contextual cues were present (e.g., neither the floors nor the walls contained any textures).

#### Morphine conditioned place preference protocol

Male Sprague-Dawley rats (12-week-old, n=12) were conditioned in a counterbalanced fashion, according to Figure 1-a. Rats were allowed free exploration of the apparatus for two days (habituation and pre-conditioning) during 15 min, and on conditioning days they received a subcutaneous injection of saline before being placed in one of the chambers for 45 min. Four hours later, animals were administered morphine (5 mg/kg SC; as in (Wu et al., 2007)) and placed in the opposite chamber for 45 min. The procedure was repeated for six days. One day after the last conditioning session, animals were tested for place preference while freely exploring the apparatus for 15 min. To measure the effectiveness of conditioning, we compared the time spent in the morphine-paired compartment by calculating the difference at post-conditioning versus pre-conditioning (CPP delta). Behavioral metrics such as time spent in compartments, distance travelled, and mean speed (without rest) were extracted from SMART video tracking system (v.3.0, Panlab, Spain). To assess differences and interactions within and between factors session and drug, we used Two-way ANOVA followed by post-hoc comparisons using Bonferroni’s correction. Values for permanence in saline- or morphine-paired compartments are reported as % time ± SEM. This protocol was validated in an independent experiment where a group of animals showed a strong preference for the drug-paired context (Supplementary figure 1-c).

**Figure 1.**
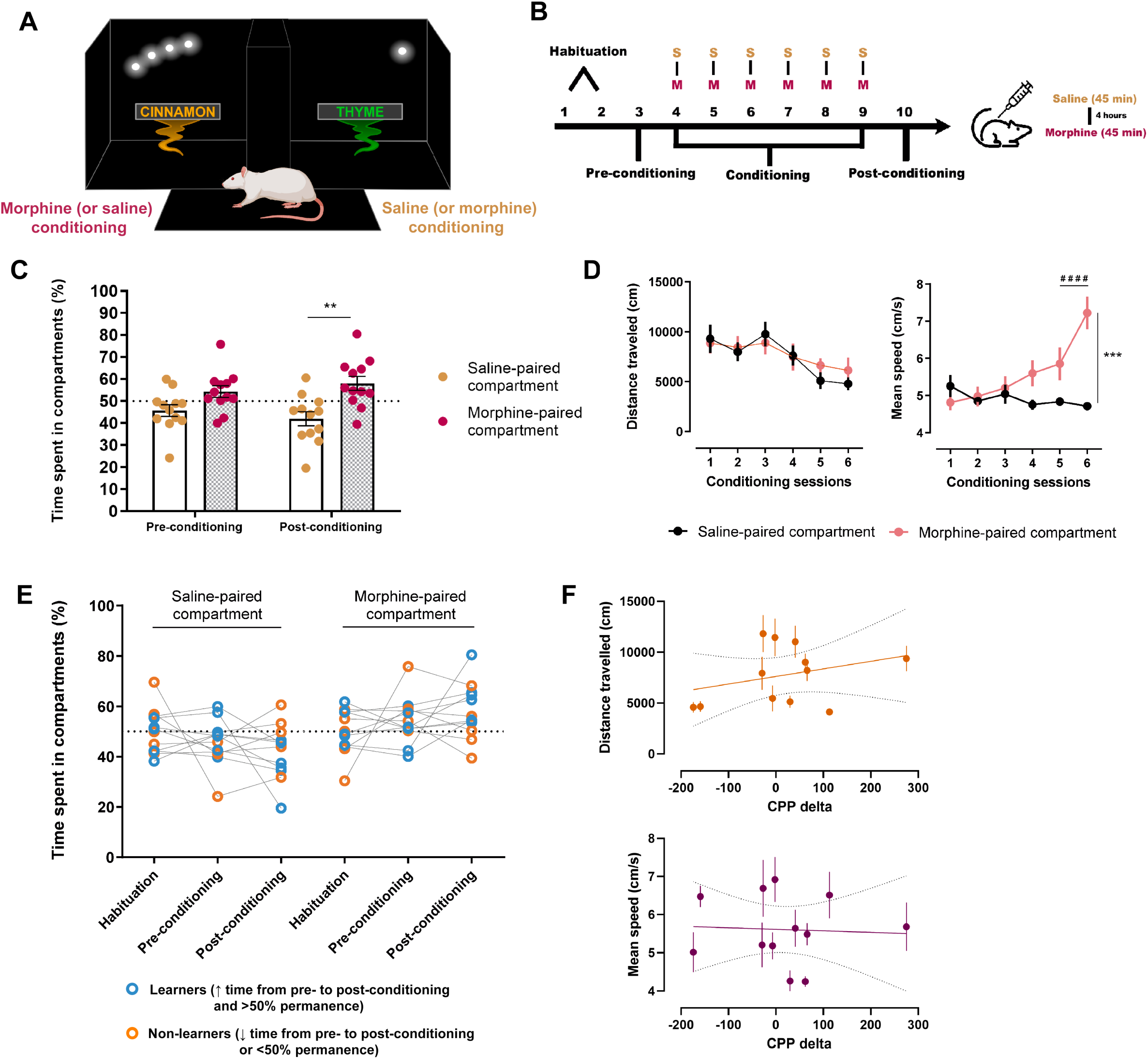
Conception of elementary and MRI-adaptable conditioned place preference contexts for functional brain imaging. a. Outline of the CPP conditioning chambers upon implementation of simple contextual cues. During free exploration of the conditioning apparatus, rats could find either a cinnamon-odorized chamber paired with small lines of white light on a dark background, or a thyme-odorized chamber illuminated by a single point of light. The unconditioned stimulus (morphine) was delivered in a counterbalanced and unbiased fashion. b. Timeline for the CPP protocol. Rats were allowed free exploration of the apparatus for two days (habituation and pre-conditioning), and on conditioning days they received a subcutaneous injection of saline before being placed in one of the chambers. Four hours later, animals were administered morphine and placed in the opposite chamber. The procedure was repeated for six days. One day after the last conditioning session, animals were tested for place preference. c. Rats developed preference for the morphine-paired context, as revealed by the percentage of time spent in the morphine-paired compartment relative to time spent in the saline-paired compartment at post-conditioning. A two-way ANOVA followed by Bonferroni’s multiple comparisons test was employed to analyze differences between times spent in the two chambers. Values are presented as mean ± SEM. **P < 0.01. d. Locomotion parameters such as distance traveled (cm) and mean speed (cm/s) were measured across conditioning sessions in both compartments. Animals showed a gradual increase in mean speed throughout conditioning in the morphine-paired compartments, while maintaining stable walking distances (two-way repeated measures ANOVA followed by Bonferroni’s multiple comparisons test). Values are presented as mean ± SEM. ####P < 0.0001; ***P = 0.0006. e. Individual permanence times in the two CPP compartments shows that M-CPP was expressed in 7 out of 12 animals. Clusters of learners and non-learners were defined based on two factors: direction of change of time spent in the compartments from pre- to post-conditioning and percentage limit (if time increased and was over 50%: learners; if time decreased or was below 50%: non-learners). f. Based on the difference between time spent in the morphine-paired compartment post- vs pre-conditioning (in sec), each animal was attributed a CPP delta value. The plots show the correlation between these delta values and the two locomotion parameters (simple linear regression, distance traveled: R^2^ = 0.094; F _(1,10)_ = 1.041, P = 0.332; mean speed: R^2^ = 0.0028; F _(1,10)_ = 0.028, P = 0.871).

### MRI experiments

We replicated the odor-visual contexts in the MRI scanner by placing optic fibers and air-delivery tubing over the MRI stereotaxic frame. MRI-compatible optical fibers (1.5 mm diameter, Chinly, China) were arranged to constitute a single or 4 linearly arranged light sources, thus reproducing the visual patterns of the CPP box. The optical fibers’ ends were placed within the focal distance of the rat’s eye, and the olfactory cues were delivered near the rat’s nose via air tubing connected to an olfactometer (Figure 2-a, left). Visual and odor stimuli were synchronized with MRI scanner trigger-out, automatically presented as pairs, synchronized using an Arduino board, thus mimicking the contextual arrangements present in the behavioral apparatus (Figure 2-a, right). We will refer to this stimulus presentation as the cue-based fMRI sequence.

**Figure 2.**
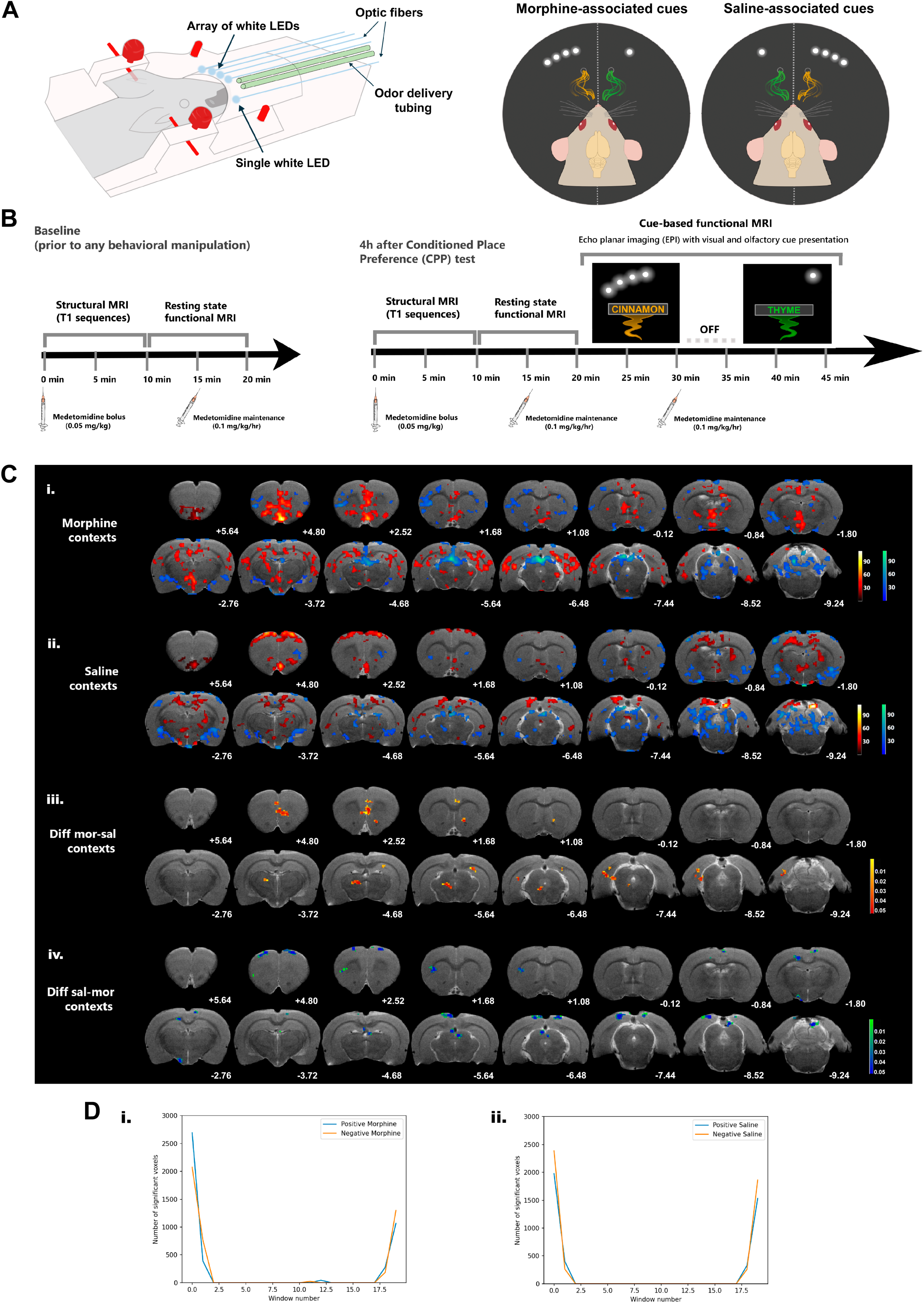
Responses to saline- and morphine-associated cues show regional segregation in the rat brain and are (re)activated early on after onset of contextual cue presentation. a. Scheme of replication of conditioning contexts in the MRI scanner. Optic fibers connected to white LEDs and air-puff tubes connected to an olfactometer were placed around the MRI stereotactic frame (left, side view; right, front view). Stimuli were presented in a counterbalanced fashion, mimicking the contexts created in the behavioral apparatus. Thus, for six animals, the morphine-paired context was cued by the array of LEDs combined with the scent of cinnamon, and the saline-paired context was cued by the single point of light combined with the scent of thyme. For the other six animals, context assignment was reversed. b. Timelines depicting our functional whole-brain imaging protocol. For all fMRI experiments, sedation was maintained by α2-adrenergic agonist medetomidine (intraperitoneal bolus at 0.05 mg/kg; subcutaneous maintenance infusion at a rate of 0.1 mg/kg/h, every 15 min after bolus injection). Animals were imaged at two time points: before and after behavioral manipulation in M-CPP. Baseline resting state functional MRI was acquired before initiation of behavioral testing (left timeline); post-CPP resting state fMRI was acquired 2 weeks after the first set of baseline acquisitions and 4h after the M-CPP test (right timeline). Anatomical scans were acquired prior to resting state functional MRI. Task-based fMRI was acquired in two 10-min blocks, and the pairs of contextual cues were presented in the same order for all animals, regardless of whether they were previously associated with morphine or saline administration in the CPP box. The two stimuli were separated by a 5 min inter-block interval. c. Representative anatomical scans (T2-weighted anatomical MRI image of our template brain) display: i. and ii. Mean BOLD activity maps within the first 30 s window in response to the morphine- and saline-associated cues, respectively. Color scale represents mean regression coefficients for morphine and saline contrasts in comparison to baseline (masked under a significance level set at 0.05 and a cluster size threshold of 16 voxels); and iii. and iv. Magnitude maps show the clusters that survived the difference between significantly activated clusters during presentation of saline and morphine cues, and vice-versa. Color scale represents P values of the surviving voxels. Coordinates under each brain section represent Paxinos & Watson’s rat brain atlas coordinates (in mm). d. Representation of the number of activated voxels in the whole brain across the 10-min context presentation sequence for the i. morphine and ii. saline contexts.

Baseline structural and resting state fMRI sequences were acquired for a few days before and right after conditioning, approximately 2 weeks within each other (according to the timeline depicted in Figure 2-a). Each animal was placed in the MRI frame on only two occasions.

Brain fMRI was carried out using a high-resolution 9.4 T small bore animal scanner (BioSpec 94/20, Bruker, Massachusetts, USA). A rat head-adapted room temperature surface coil was combined with a volume transmission coil, and acquisitions were made using Paravision 6.0.1 software (PV6, Bruker). Anatomical images were acquired using turbo rapid acquisition with relaxation enhancement (Turbo-RARE) T2-weighted sequence (22 continuous slices with 0.9 mm, TE=33.00 ms, TR 2.5 sec, resolution=0.098*0.098 mm and averages=2), as a standard procedure in rodent structural imaging, allowing us to keep an adequate acquisition time at high-field. Using Free Induction Decay echo-planar imaging (FID-EPI) and a repetition time (TR) of 2 sec, fMRI data were acquired in 2 blocks for the post-behavior imaging session, 30 min after medetomidine and isoflurane clearance: resting state-fMRI for 10 minutes, with a 2-min interval before initiating the cue-based fMRI. To monitor brain activity during contextual memory recall, we targeted the low frequency dynamics of BOLD signal for an extended temporal window, thus dividing visual-olfactory cue presentation into two continuous 10-min blocks, to mimic a time frame of free exploration, memory reactivation, and conditioned response, to which animals were exposed in the CPP apparatus on the post-conditioning (or test) day. For all animals, the first 10-min block presented corresponded to the “cinnamon + white light 4 LED array”, and the second to the “thyme + white light point” contexts. Both contexts were separated by a 5-min inter-block pause. Timelines for the fMRI experiments can be found in Figure 2-b. Although the two contexts were presented in the same order, the associated outcome (context valence) counterbalanced the context presentation order. Consequently, six rats first encountered the morphine-paired cues, and the other six were first exposed to the saline-paired cues. Sixteen axial slices of 0.9 mm thickness were recorded to cover the main areas of interest in the brain (echo time/repetition time = 15/2000 msec, spatial resolution = 0.234 * 0.234 * 0.9, matrix 128 * 128, field of view (FOV) 30 * 30 mm, bandwidth=400000 Hz). After imaging procedures, animals received a subcutaneous bolus injection of 0.1 mg/kg atipamezole (Antisedan, Pfizer, Karlsruhe, Germany), to counteract the effects of medetomidine, and were placed in a recovery box for post-scan monitoring until they were fully awake.

### fMRI data preprocessing and analysis

Imaging data were preprocessed using SPM12 (statistical parametrical mapping) toolbox for MATLAB (R2019b). Preprocessing steps included realignment of images to the first volume (or scan) of each acquired sequence, normalization to a common rat brain template (affine transformation) to account for individual anatomical and positional differences, and smoothing for signal-to-noise ratio improvement (Gaussian kernel with FWHM of twice voxel size). A final filtering step using a 0.01-0.1 Hz band-pass filter was performed using the RESTplus toolbox (v1.25_20210630) for MATLAB to extract low frequency fluctuations specific to the BOLD signal.

For activity analysis, we fit a single generalized linear model (GLM), implemented in Python using Nipy (Gorgolewski et al., 2011), to our continuous BOLD data by concatenating the recordings of all 12 rats accounting for the context presentation order. The two 10-min blocks were partitioned into 30-sec windows (total = 40 windows) to be sensitive to changes in response over the long duration of contextual cue presentation. A regressor was added to the design matrix for each window using a rat adjusted hemodynamic response function (Lambers et al., 2020). The 6 realignment parameters and the whole brain mean signal of individual rats were also added as nuisance regressors. We considered contrasts for individual windows and conditions (contexts) in addition to differences between conditions for matched windows (e.g. first window saline minus first window morphine). All comparisons were FDR-corrected with a significance level set at 0.05 and a cluster size threshold of 16 voxels (0.79 mm^3^).

For functional connectivity (FC) analysis, we selected some regions for seed-based and region-of-interest (ROI)-based static connectivity analysis. Seed regions were selected from the cortico-limbic system involved in memory processing (encoding and retrieval), affective state, and valence/salience assignment, including three amygdala nuclei (anterior cortical amygdaloid nucleus, ACo, basomedial BMA, and central, CeA, amygdala), the intermediate portion of the lateral septum (LSI), lateral habenula (LHb), ventral tegmental area (VTA), two hippocampal subregions (dentate gyrus, DG, and CA1 subfield), and two cortical structures, cingulate (Cg) and retrosplenial (RSC) cortices. ROIs were manually defined by drawing masks of varying voxel size over these regions bilaterally across the template brain images using MRIcron (RRID:SCR_002403) software. Depending on the region’s antero-posterior extension, some masks spanned across multiple slices and were combined into a single mask. Adobe Photoshop (CC 2019) was used to confirm correct mask positioning by superimposing Paxinos & Watson’s rat brain atlas figures (Paxinos & Watson, 2007) over the sixteen template functional brain slices covering the whole brain.

For ROI-based analysis, a MATLAB-written script was developed to extract timeseries data from individual ROIs as well as pairwise Pearson’s correlation coefficients from all possible pairs of ROIs. FC matrices were z-transformed (Fisher’s z-score) and obtained for baseline (pre-CPP) and post-CPP resting state sequences. Correlation values were compared at the ROI-pair level between the two conditions using paired t-tests with false discovery rate (FDR) corrections. P-value matrices (uncorrected and false discovery rate (FDR)-corrected) displaying statistically significant differences between the two acquisition moments were also computed. A separate set of matrices was extracted from the ‘context recalling’ sequence and represented as correlation matrices of functional connectivity for conditions ‘morphine’ and ‘saline’. To ensure that animals perceived contextual cue presentation under (light) sedation, we also performed pairwise comparisons between resting state and cue state sequences (pre-CPP and post-CPP resting state individually compared to morphine- and saline-paired contexts). FC matrices were also extracted for other limbic and sensory information processing brain regions (Supplementary data - figure S4).

For seed-based analysis, the same user-defined ROI masks were used as seeds for voxel wise whole brain connectivity analysis. The RESTplus toolbox retrieved zFC maps for each subject which were later combined into group mean maps for each of the four study conditions. Pairwise paired t-tests were performed on these images, and t maps were obtained and corrected for multiple comparisons using the Gaussian Random Field Theory procedure (one-tailed with voxel- and cluster-level p values set at 0.05 were used as statistical map correction parameters). Positive voxel t values indicated first condition greater than second; for resting state analysis, post-CPP corresponds to condition 1 and pre-CPP to condition 2. For cue-based analysis, ‘morphine context’ represents condition 1 and ‘saline context’ condition 2. In the end statistical maps were overlaid onto template anatomical T1 brain images. An in-house MATLAB script was used to extract mean functional connectivity data from each seed to measure their intra-network relevance across the four study conditions.

Graph theoretical representations of the cortico-limbic network under study were extracted from the MATLAB-based visualization toolbox BrainNet Viewer (v.1.7, Xia et al., 2013). Node files containing ROI mask x-y-z coordinates, node centrality degree, and its correspondent color coding were created. For graph representation, we extracted only unilateral coordinates of our ROI masks from MRIcron. Node centrality degree was calculated as the sum of all connections each node (ROI mask) had in our undirected network, based on the work by Centeno et al. (2022). Node color was defined from a direct conversion of nodal degree into the jet colormap code in MATLAB (scale limits were set based on the least and most central nodes in our analysis). Edge files contained the correlation matrices extracted from our ROI-based analysis. Purely for visualization purposes, edge threshold was arbitrarily set at a z-score of 0.2. For the surface file, we used the SIGMA Rat Functional Imaging EPI Brain Template (Barrière et al., 2019).

To assess whether baseline brain functional connectivity predicts performance in the CPP test, we performed a simple linear regression and plotted the CPP delta against the ROI-based correlation coefficients of all possible ROI pairs for resting state conditions ‘pre-CPP’ and ‘post-CPP’. Pearson’s R was computed along with significance values. We conducted the same type of analysis for ‘context recalling’ conditions under the hypothesis that performance in the CPP test could be correlated to pairwise connectivity strength during contextual cue exposure. Our general prediction was that subjects scoring higher in the CPP test would have a higher functional connectivity in memory-related and/or valence assignment regions during presentation of morphine-paired contexts compared to the level of connectivity during exposure to saline-paired contexts. For rats that scored low or negative deltas, either the reverse or the absence of a correlational effect would be observed.

## RESULTS

### Validation of a new MRI-compatible conditioned place preference protocol

Animals displayed a preference for the morphine-paired compartment during post-conditioning (Figure 1-c, two-way ANOVA, no session x compartment interaction; compartment effect: F_(1,44)_ = 17.94, P = 0.0001; Bonferroni-corrected multiple comparisons and mean % ± SEM: post-conditioning saline (41.96 ± 3.14) x post-conditioning morphine (58.04 ± 3.14): P = 0.0019). No significant change was observed for morphine-paired compartment preference between pre- and post-conditioning. Throughout conditioning, morphine administration did not impact locomotor activity as measured by the cumulative walking distance in the enclosed compartments (two-way repeated measures ANOVA, no session x compartment interaction: F _(5, 110)_ = 0.77, P = 0.574), but a strong effect was found on the mean speed (without considering rest time) as it gradually increased and significantly peaked on the last conditioning day (two-way repeated measures ANOVA, session x compartment: F _(5, 110)_ = 12.93, P < 0.0001; Bonferroni-corrected multiple comparisons and mean (cm/s) ± SEM: day 6 saline-paired compartment (4.712 ± 0.114) x day 6 morphine-paired compartment (7.224 ± 0.437); P = 0.0006); day 5 morphine-paired compartment (5.853 ± 0.437) x day 6 morphine-paired compartment (7.224 ± 0.437); P < 0.0001). Those rats showing both an increased time spent in the morphine-paired compartment from pre- to post-conditioning and a higher than 50% permanence therein were classified as “learners” (7/12) (Figure 1-e blue dots). Five of the twelve rats showed a 7% to 27% decrease in time spent in the morphine-paired compartment in the test phase and were considered “non-learners” as a result (Figure 1-e, orange dots).

By computing the difference between the time spent in the morphine-paired compartment post- vs pre-conditioning (in sec) we can attribute to each individual animal a CPP delta value reflecting how effective was the association between context and morphine. We did not find a correlation between CPP delta and the two locomotion parameters evaluated in the present study (Figure 1-f top; distance traveled: R^2^ = 0.094; F _(1,10)_ = 1.041, P = 0.332; Figure 1-f bottom; mean speed: R^2^ = 0.003; F _(1,10)_ = 0.028, P = 0.871), indicating that our morphine-induced place preference did not result from the locomotor reactivity to morphine.

### Morphine- and Saline-associated contextual stimuli reactivate common, and specific, hippocampal-prefrontal-mesolimbic areas around stimulus onset

Once analyzed using the GLM model, MRI activity maps revealed both context-specific and context-general activity clusters across the whole brain in the first 30 s of cue presentation (Figure 2-c). Morphine-associated stimuli presentation (Figure 2-ci) resulted in increased activity in prefrontal regions, namely prelimbic (PrL), infralimbic (IL), orbitofrontal cortices (medial - MO, ventral - VO, and lateral - LO), hippocampal and parahipocampal regions, namely CA2 and CA3, dorsal subiculum (DS), dorsolateral entorhinal (DLEnt), medial entorhinal (MEnt), and perirhinal (PRh) cortices, connected subcortical areas bed nucleus of the stria terminalis (BNST), anteromedial (AM), lateral and medial geniculate (MG) thalamic nuclei, parvicellular region of the red nucleus (RPC), the pontine reticular nucleus (PnO), posterior hypothalamic area (PHA), anterior pretectal nucleus (APT), and VTA parainterfascicular (PIF) nucleus. Saline-associated stimuli increased activity over the antero-posterior (AP) motor cortex, the forelimb region of primary somatosensory cortex (S1FL), medial septum (MS), dysgranular retrosplenial cortex (RSD), dorsomedial periaqueductal gray (DMPAG), ventral CA1, and dorsal cortex inferior colliculus (DCIC) (Figure 2-cii). Activation of some of these regions has been shown in response to withdrawal-related cues in the rat, namely the thalamic and hypothalamic regions (Carmack et al., 2019), but at least the activation of cognitive and memory regions seems to be specific to morphine memory retrieval, rather than responses to withdrawal states.

Context-general activated regions tended to elicit stronger activity in response to morphine-associated stimuli. These included the cingulate cortex (Cg), the core (AcbC) and shell (AcbSh) subdivisions of the nucleus accumbens, and primary (Au1) and secondary (AuD and AuV) auditory cortices. Very similar levels of activity were found in the anterior olfactory nuclei (AOL, AOV), and its cortical output piriform cortex (Pir), primary (V1) and secondary (V2) visual cortices, intermediate portion of the lateral septum (LSI), dorsal caudate putamen (CPu), the hindlimb (S1HL) and trunk (S1Tr) regions of the primary somatosensory cortex, medial preoptic nucleus (MPO) and area (MPA), nucleus reuniens (Re), ventromedial (VMH) and posterior (PH) hypothalamus, septo-hypothalamic nucleus (SHy), hippocampal CA1 and DG subregions, rostral portion (VTAR) and parabrachial pigmented nucleus (PBP) of VTA, medial (MPtA) and lateral (LPtA) parietal association cortices, ectorhinal cortex (Ect), and lastly the temporal associative cortex (TeA). Most of the context-general activation is bilateral. Exceptions, unilateral and in opposite hemispheres, are VTAR and PBP (morphine-associated cues – left hemisphere; saline-associated cues – right hemisphere).

Just like positive clusters, we found negative clusters throughout the whole brain and mostly located over the insular cortex, somatosensory cortex, amygdala subregions, ventral hippocampus, and superior colliculus in both conditions.

We then compared activity, still in the first 30-s window, in identified regions between morphine and saline-associated contexts by looking at the signal magnitude differences (Figures 2-Ciii; Civ). Significant clusters of increased activity in morphine relative to saline contexts were found over orbitofrontal and medial pre-frontal PrL and IL cortices, anterior Cg (anterior), posterior RSD, AcbC, post-thalamic nuclear group (Po), CA1, APT, p1RF, DS, posterior DG, presubiculum (PrS). Conversely, we found clusters with higher activity in saline relative to morphine contexts over the motor cortex, primary somatosensory cortex, CPu, Cg (posterior), tuberal region of the lateral hypothalamus, RSC, distal CA1, medial pretectal nucleus (MPT), primary and secondary visual cortices, retrosplenial dysgranular cortex, and superior and inferior colliculi. Altogether these results demonstrate that saline-associated cues activate regions involved in somatosensory processing, while morphine-associated cues activate higher cognition, limbic, and memory regions.

Significantly increased and decreased activity clusters appear primarily in the first 30-sec window, and only again in the last two acquisition windows (Figure 2-d, Supplementary figure 2). That was the case for several subdivisions of the insular cortex (granular, agranular, and dysgranular), amygdala (central, basolateral and basomedial) and amygdaloid nuclei (anterior cortical, anterior area, lateral and medial amygdaloid), ventral pallidus, ventral claustrum, ventral subiculum, dorsal raphe, lateral parabrachial nucleus, superior colliculus, premammillary nucleus, substantia nigra, and ventral DG. Specific to saline contexts (Figure 2-dii) were VO and LO, DS, medial pretectal nucleus, AcbC, granular and dysgranular insular cortex, substantia nigra, ventral CA1, ventral DG, and MEnt, which had not appeared active in the first window. Specific to morphine contexts (Figure 2-di) we found ACo, LPtA, MPtA, ventral pallidus, ventral claustrum, pyramidal cell layer of anterior CA1, dorsal raphe, LPAG, and lateral parabrachial nucleus.

### Drug-context pairings promoted low-level rearrangements of resting state cortico- limbic network connectivity

We hypothesized that morphine-context pairings during CPP would impact neural processing of sensory, mnemonic, and emotional information, revealed by overall changes in correlated BOLD signal across pertinent neural populations. We thus focused on pair-wise comparisons of resting-state BOLD signal amongst ten empirically selected regions of a putative cortico-limbic network, before and after behavioral manipulations. In both pre- and post-CPP our ROI-based analysis revealed significantly correlated activity across CA1, DG, Cg, and RSC, regions involved in spatial contextual memory processing and consolidation (Figure 3-a top, z-scored matrix). This is consistent with these regions’ well-known engagement in the default-mode network, which is involved in episodic memory (Lu et al., 2012). In addition, we found that the two amygdala nuclei were correlated with each other and with one of their cortical counterparts, ACo, in both condition***s.*** When we compared the set of correlation coefficients for each ROI pair between pre- and post-CPP, and found a significant difference between the pre- vs post-connectivity in the lateral habenula (LHb)-hippocampal CA1 pair (Figure 3-a bottom, paired t-test, pre-x post-CPP Pearson’s correlation coefficients, FDR-corrected P = 0.014).

**Figure 3.**
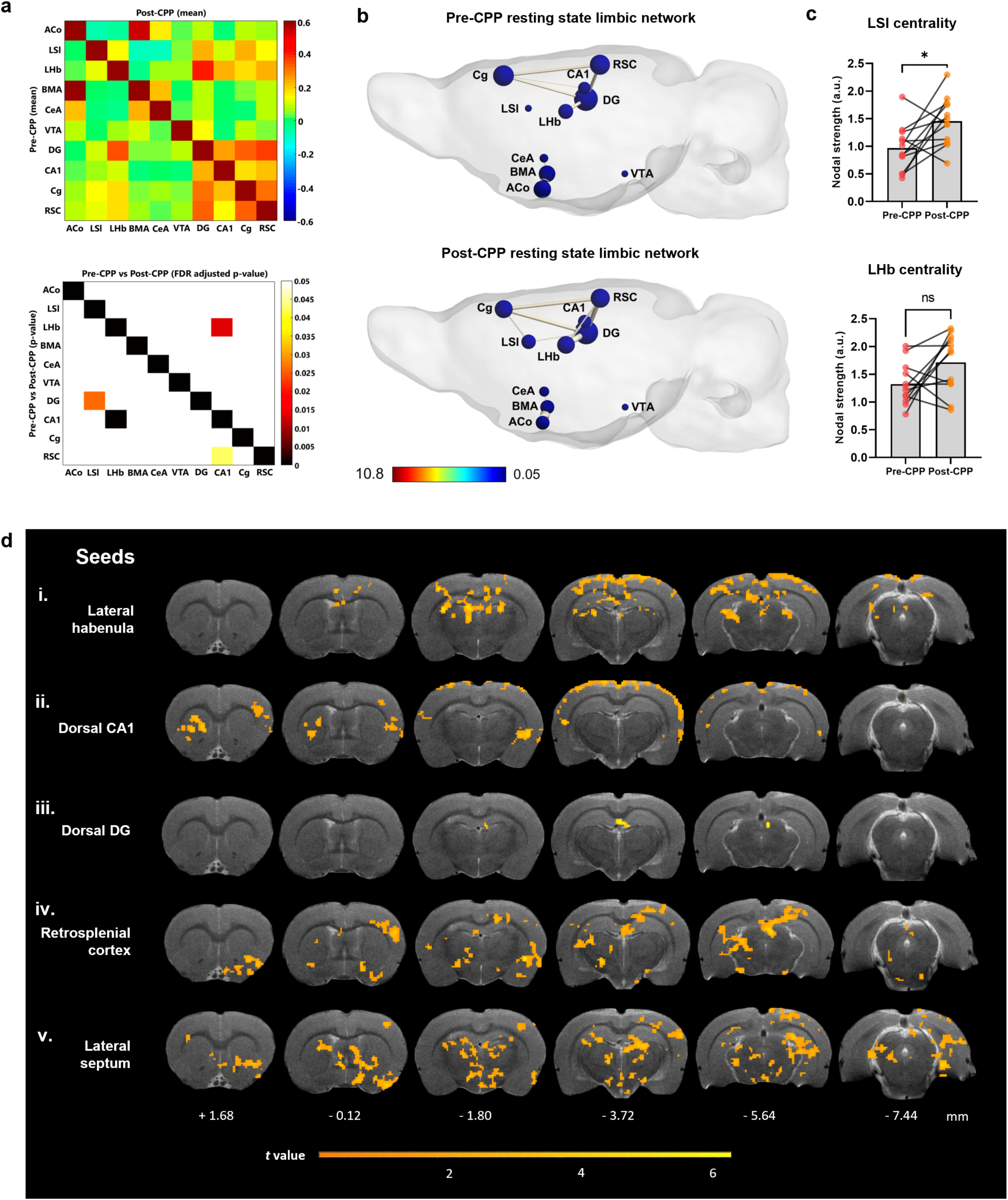
Drug-context pairings promoted low-level rearrangements of resting state cortico-limbic network connectivity. a. Functional connectivity (FC) increases after morphine conditioning. Upper panel: Mean FC-matrices of pre-CPP (lower half) and post-CPP (upper half). X and Y-axes represent brain regions. Color scale represents Pearson correlations i.e., strength of FC between each pair of brain regions. Lower panel: p value matrix (lower half) representing statistically significant differences between pre-CPP versus post-CPP resting state connectivity. FDR-adjusted p value representing statistically significant differences between pre-CPP versus post-CPP resting state connectivity (upper half). Color scale represents p values (0-0.05). X- and Y-axes represent brain regions. Abbreviations: ACo = anterior cortical amygdaloid nucleus, LSI = lateral septum intermediate, LHb = lateral habenula, BMA = basomedial amygdala, CeA = central amygdala, VTA = ventral tegmental area, DG = dentate gyrus, CA1 = CA1 region of the hippocampus, Cg = cingulate cortex, RSC = retrosplenial cortex. b. Graph theory representation of the putative cortico-limbic network in pre- and post-conditioning at rest. Color scale represents node color, which was defined from a direct conversion of nodal degree into the jet colormap code in MATLAB (scale limits were set based on the least and most central nodes in our analysis). c. Plots depict centrality levels of two nodes of the network, the LSI and the LHb. Values are presented as mean nodal strength ± SEM. *P < 0.05 (paired t test). d. Representative anatomical scans (T2-weighted anatomical MRI image of our template brain) display mean statistical t maps of resting state connectivity for five regions within our putative cortico-limbic network, converted into seeds for seed-based analysis. Color scale represents t-value i.e., strength of functional connectivity of the seed region with all voxels in the brain. Coordinates under each selected brain section represent Paxinos & Watson’s rat brain atlas coordinates (in mm).

We then looked at network-level interactions by using graph theoretical analysis, and analyzing nodes and edges as proxies for brain region activity levels and functional connections between region pairs, respectively (Sporns, 2018). Nodal degree, also known as centrality, reflects the connection density of each individual brain region, e.g. its importance in a given functional network (Centeno et al., 2022). Here we measured centrality as the sum of all correlation coefficients of an individual region with every other region in the proposed network, and found that the LSI exhibits a significant increase in centrality from pre- to post-CPP (Figure 3-b bottom, 3-c top, two-tailed paired t-test, t = 2.735, df=11, P = 0.019), which was not the case for LHb (Figure 3-c bottom, two-tailed paired t-test, t = 2.069, df=11, P = 0.063). This suggests that both positive and negative affect processes, engaging episodic memory and reward (or withdrawal)-related regions, may occur throughout the morphine conditioning protocol, in alignment with the acute shifts between rewarding and aversive states during the development of drug addiction.

To expand our analysis beyond pair-wise correlations in cortico-limbic circuits, we selected the five regions whose connectivity changed from pre- to post-CPP and used them to seed voxel-wise whole-brain connectivity analysis: LSI, LHb, CA1, DG, and RSC. Statistical maps in Figure 3 show, for each seed region, Gaussian-corrected one-tailed t-maps for post-CPP vs pre-CPP comparisons (with a voxel and cluster threshold set at 0.05), highlighting the regions whose activity exhibited higher correlation in the post-CPP resting state versus pre-CPP (Figure 3-d).

In this analysis, the lateral habenula (LHb), a region involved in aversive learning, is highly correlated with medial mesocortical cingulate and retrosplenial areas, and with visual, somatosensory, medial, and lateral parietal regions, and dorsal CA1, in line with the increase in resting state connectivity observed in the ROI-based analysis explained above (Figure 3-di). Dorsal CA1 was significantly more correlated with striatal CPu, SS, RSC, temporal associative and ectorhinal areas, but not LHb (Figure 3-dii). Hippocampal DG showed two evident clusters, one over the fasciola cinereum (FC), distal CA1, and DG itself, and the other over the superior colliculus (Figure 3-diii). RSC exhibited increased post-CPP correlations with widespread brain areas, namely: hippocampal dSUB, CA1, DG, central and basal amygdala, insula, claustrum, substantia nigra, accumbens, superior colliculus, parietal cortices, posterior hypothalamus, and lateral olfactory tract nucleus, regions involved in ego and allocentric contextual encoding, memory, and affective processing (Figure 3-div). Thus, following CPP there seems to be a dissociated emergence of two, known, distinct networks, an affective one centered in the LHb, and a spatial-contextual-mnemonic one centered around HIPP-RSC.

Notably, the lateral septum, the strongest non-hippocampal target of hippocampal CA3 outputs, and a significant source of inputs to VTA, exhibited increased post-CPP correlations with the highest number of brain areas, consistent with the increases in centrality observed in the ROI-based analysis explained above in post-CPP resting state (Figure 3-dv). Here it is noteworthy the connectivity with regions of the multi-sensory integration and spatial contextual processing network (*hippocampus, presubiculum, orbitofrontal, insular, piriform, somatosensory, parietal and temporal association, visual, auditory, retrosplenial, and medial entorhinal cortices*), and with affective and sensory processing-related regions such as nucleus accumbens core, hypothalamic (*e.g. lateral preoptic area*) and thalamic nuclei (*e.g. nucleus reuniens and medial geniculate nucleus*), BNST, LHb, PAG, and VTA. In this case, LSI seems to emerge as the central hub of a functional network linking the two dissociated emergent networks, affective and spatial-contextual-mnemonic, discussed above.

*Other seed region connectivity can be found as supplementary data (Supplementary figure 3)*.

### A common limbic circuit (likely) relies on affective state to retrieve contexts with opposed value upon presentation of saline or morphine-associated cues

Why are there two functionally distinct networks emerging independently upon morphine-induced CPP, and linked by LSI?

The olfactory and visual systems, responsible for processing and integrating sensory information, were expected to be highly engaged during presentation of the two contextual cue modalities. We investigated their intra-system correlation coefficients in the four study conditions (pre- and post-CPP resting state, and saline- and morphine-associated cues presentation) and found a significant difference between the pre- vs post-CPP connectivity in the anterior olfactory nucleus (AON)-nucleus of the lateral olfactory tract (LOT) pair (Supplementary Figure 4, paired t-test, pre- x post-CPP Pearson’s correlation coefficients, FDR-corrected P = 0.047). No further differences were found between pre- vs post-CPP and saline- vs morphine-associated cue connectivity in olfactory and visual regions. We then compared their coactivity with cognitive/emotional regions by randomly selecting four cortico-limbic network regions (see above) and two entorhinal cortex regions, which are well known gateways for sensory information entering in the hippocampus (Canto et al., 2008; Eichenbaum & Lipton, 2008). Here we found that presentation of morphine-associated cues elicited a significant change in the patterns of coactivity when compared to post-CPP resting state connectivity. Most strikingly, connectivity within visual regions, and their connectivity with CA1 and DG were both significantly altered (Supplementary Figure 4-fi, morphine vs post-CPP matrix), which was not the case for olfactory regions (Supplementary figure 4-ci, morphine vs post-CPP matrix), suggesting that visual contextual sensory encoding dominates the initial acquisition of a contextual representation, and likewise its subsequent association with affective stimuli.

Having found an activity network whose connectivity stores the representation of CPP, we then asked whether such network specifically retrieves distinct contextual representations when the animal is presented with the sensory cues that define each opposed valence context. For this, we analyzed the synchronicity across such network during a 10 min presentation of either saline or morphine-associated cues. In agreement with the data shown above, we observed strong correlations between spatial contextual and episodic memory processing regions (Figure 4-a top, z-score matrix, DG, CA1, Cg, and RSC), and between these and the LSI, LHb, and VTA, with no significant differences between the two contexts (Figure 4-a bottom, p value matrix).

**Figure 4.**
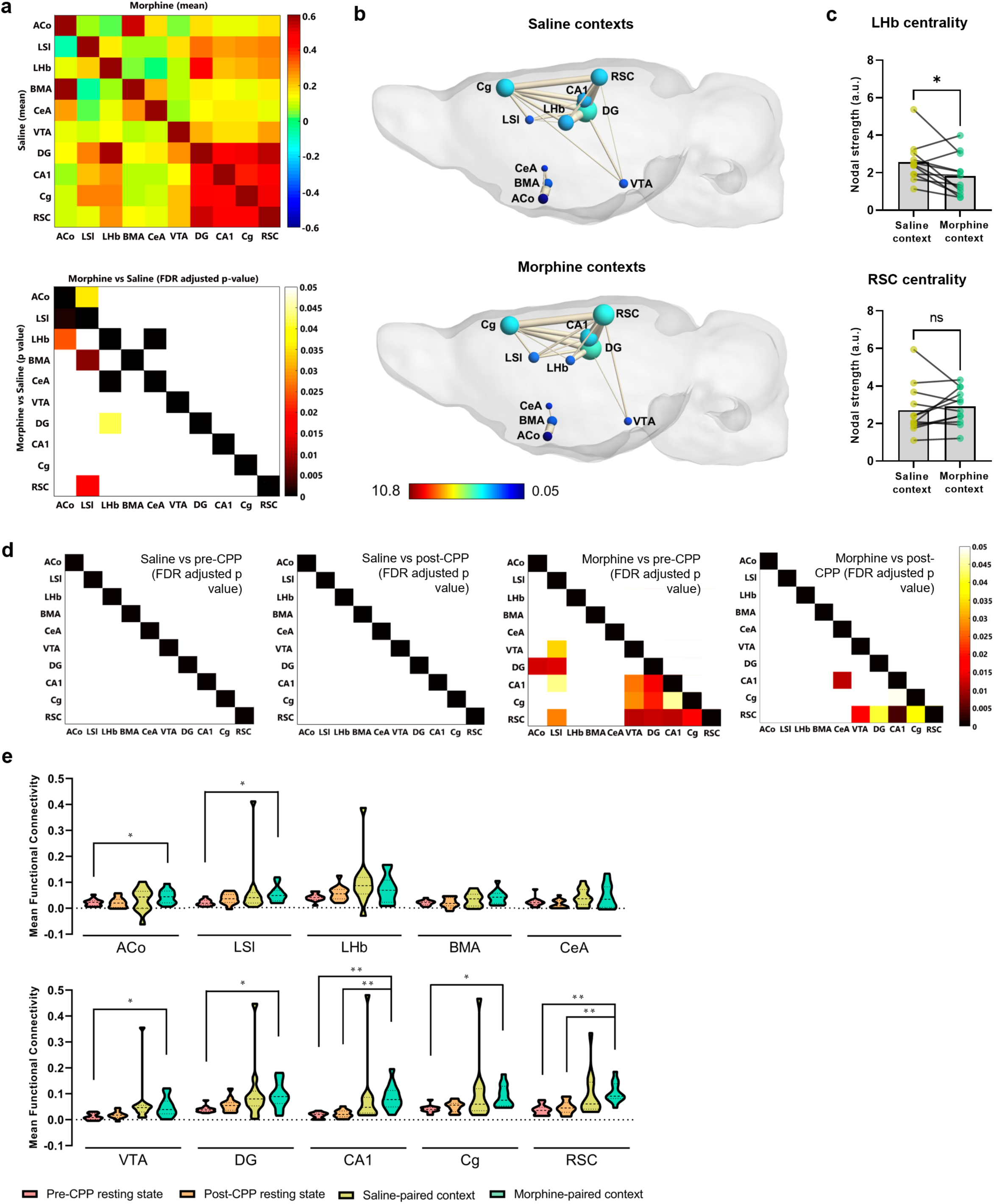
The same limbic circuits are involved in the retrieval of distinctly valenced contexts, but discrimination likely depends on affective state. a. Functional connectivity (FC) between saline- and morphine-paired contexts. Upper panel: Mean FC-matrices of saline (lower half) and morphine (upper half) contexts. X and Y-axes represent brain regions. Color scale represents Pearson correlations i.e., strength of FC between each pair of brain regions. Lower panel: p value matrix (lower half) representing statistically significant connectivity differences between saline versus morphine contexts. FDR-adjusted p value representing statistically significant connectivity differences between saline versus morphine (upper half). Color scale represents p values (0-0.05). X- and Y-axes represent brain regions. Abbreviations: ACo = anterior cortical amygdaloid nucleus, LSI = lateral septum intermediate, LHb = lateral habenula, BMA = basomedial amygdala, CeA = central amygdala, VTA = ventral tegmental area, DG = dentate gyrus, CA1 = CA1 region of the hippocampus, Cg = cingulate cortex, RSC = retrosplenial cortex. b. Graph theory representation of the putative cortico-limbic network during exposure to saline and morphine contexts. Color scale represents node color, which was defined from a direct conversion of nodal degree into the jet colormap code in MATLAB (scale limits were set based on the least and most central nodes in our analysis). c. Plots depict centrality levels of two nodes of the network, the LHb and the RSC. Values are presented as mean nodal strength ± SEM. *P < 0.05 (paired t test). d. FDR-adjusted p value matrices representing statistically significant connectivity differences saline context exposure and the two resting state conditions (two left matrices), and morphine context exposure and the two resting state conditions (two right matrices). Color scale represents p values (0-0.05). e. Mean FC plots representing mean FC of the 10 seed regions with other voxels in the brain for all fMRI conditions. *P < 0.05, **P < 0.01 (repeated measures one-way ANOVA).

Neural activity in two amygdala regions was negatively correlated with that in either LSI or LHb when animals were presented with sensory cues defining opposed valence contexts. Namely, presentation of saline contextual cues resulted in a negative correlation between the cortical amygdaloid nucleus and the intermediate lateral septum, whereas in the morphine case a negative correlation was found between the central nucleus of the amygdala and the lateral habenula (Figure 4-a bottom, paired t-test, FDR-corrected z = -0.057, P = 0.036; z = -0.013, P = 0.00068, respectively). We then analyzed, as before, the average centrality of each of the candidate regions during contextual presentations. We found that exposure to morphine-associated cues globally decreased centrality in the lateral habenula, when compared to the one obtained during saline-associated context exposure (Figure 4-b,c; two-tailed paired t-test, t = 2.566, df=11, P = 0.026). Of all candidate regions analyzed, the retrosplenial cortex (RSC), highly connected with a cortico-limbic network involved in ego and allocentric contextual encoding, exhibits the highest increases in correlations with most of this network on exposure to morphine-associated cues relative to resting state conditions (Figure 4-d two right matrices), but no increases in centrality when compared with saline-associated cues (Figure 4-c bottom, two-tailed paired t-test, t = 0.716, df=11, P = 0.489). Interestingly, network connectivity on exposure to saline-associated cues did not differ from that observed in resting state conditions (Figure 4-d two left matrices). This reflects a possible role of the RSC in storing CPP cues of opposed valence, while discrimination may rely on inputs from or outputs to regions more dedicated to affective processing, like the lateral septum or the lateral habenula.

Several other regions exhibited increased functional interconnectivity when presented with morphine-associated contexts relative to resting state conditions, namely the lateral septum, hippocampal DG and CA1, Cg, and VTA (Figure 4-d, two right matrices). Such enhanced recruitment of spatial-contextual memory regions accompanied by increased connectivity with LSI strongly suggests that the above-mentioned two dissociated emerging networks, affective and spatial-contextual memory, fulfill distinct functions, linked by the mutually connected lateral septum. Furthermore, morphine-context retrieval elicited significant increases in brain-wide connectivity for most of the ten seed regions in our network of interest when compared to the pre-CPP resting state condition, except for LHb, BMA, and CeA. CA1 and RSC exhibited an additional difference with the post-CPP resting state condition (Figure 4-e; repeated measures one-way ANOVA, followed by Bonferroni’s multiple comparisons corrections, *p* value threshold set at 0.05).

### Pre-CPP resting state connectivity significantly correlates with behavioral CPP-delta

We found a considerable level of variability in CPP scores in our group of rats that was not directly linked to a locomotor response to morphine. We thus hypothesized that this variation in the development of CPP reflected individual intrinsic connectivity of the neural circuitry analyzed above. To test this hypothesis, we performed simple linear regression on every possible pair of regions within our putative cortico-limbic network and plotted the z-scored connectivity for each resting state condition against the individual CPP deltas. Of the 45 connected pairs hypothesized, we found a significant regression in 7 pairs, including the regions found to be changed before: LHb, CeA, LSI, and ACo. Four of these pairs included the lateral habenula, a region seen before (above) to exhibit increased resting state connectivity and decreased centrality during morphine context cue presentation, and a negative correlation with the amygdala (Figure 5-a).

**Figure 5.**
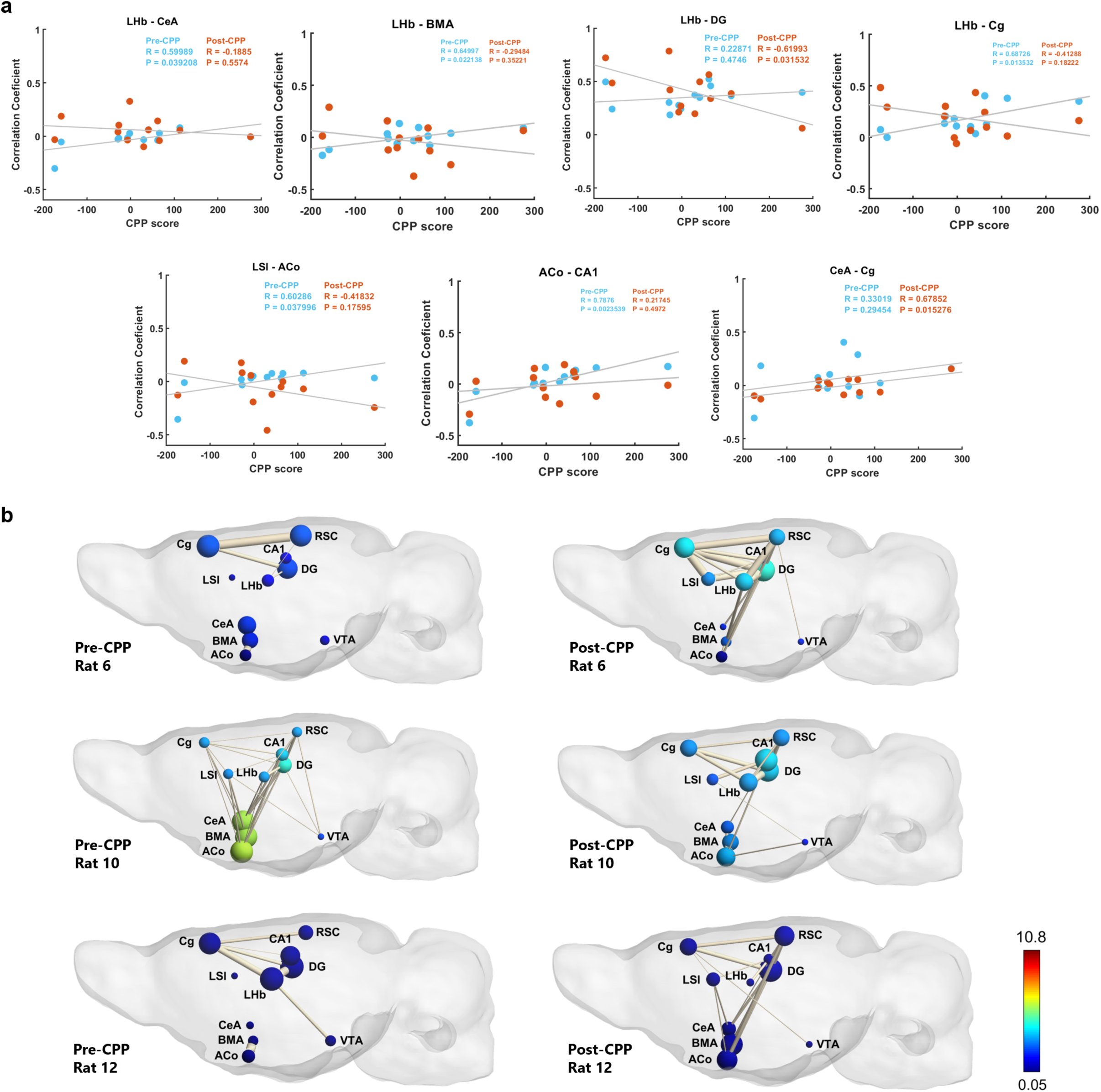
Resting state habenular and amygdala subcircuitry predicts scoring levels in the morphine-CPP test. a. Statistically significant correlation plots between resting state connectivity in the indicated regions of the network and CPP delta. Values represent the CPP delta and the correlation coefficient for each individual animal in pre-CPP (blue) and post-CPP (orange) conditions. R^2^ and p values are indicated in each panel. b. Graph theory representation of the putative cortico-limbic network in pre- and post-conditioning at rest for three border animals, two bad (rats 6 and 10) and one good learner (rat 12). Color scale represents node color, which was defined from a direct conversion of nodal degree into the jet colormap code in MATLAB (scale limits were set based on the least and most central nodes in our analysis).

Resting state connectivity before CPP was found to be significantly correlated with CPP-delta, indicating that some animals are more prone to morphine conditioning. This correlation was positive in the 7 region pairs, but only significant in 5: LHb-CeA, LHb-BMA, LHb-Cg, LSI-ACo, and ACo-CA1. This result is even stronger than the connectivity after CPP. Here, except in the case of LHb-DG and CeA-Cg pairs, all other regions showed weaker post-CPP regressions, usually accompanied by a negative regression slope and increased data dispersion. Animals with increased post-CPP connectivity usually exhibited a lower behavioral CPP-delta, and vice-versa. The fact that pre-CPP connectivity, namely within the habenula and amygdala regions, is the best predictor of future CPP-delta, with post-CPP connectivity correlated with behavioral performance only for two pairs of regions (LHb-DG and CeA-Cg), strongly suggests that some animals are more pre-disposed to CPP by virtue of their individual neural circuit architecture and perhaps plasticity states. This raises the interesting possibility that some animals have a higher susceptibility for drug addiction, by virtue of their brains being somehow richer in its neural determinants.

The above point is further illustrated if we focus on 3 animals of the group (Figure 5-b). We refer to them as border animals, as they scored highest (rat 12, CPP delta = 275) and lowest (rat 6, CPP delta = -159; rat 10, CPP delta = -174) in the CPP test. Their individual graph theoretical representations of putative cortico-limbic network activity reveal distinct connectivity patterns. Centrality of amygdala subregions in pre-CPP resting state is observably higher in the two lowest scoring animals than in the highest scoring animal. The post-CPP resting state amygdala centrality slightly changed, and we observed a higher correlation of amygdala and episodic memory regions in the latter rat. LHb centrality was also visibly different between the highest and lowest scoring animals, having respectively decreased and increased its degree from pre- to post-CPP. Episodic memory-related regions did not seem to show a marked difference in their centralities between animals and resting state conditions, suggesting that CPP development, although dependent on two networks, one spatial-contextual-mnemonic and another affective, might depend on intrinsic connectivity among affective/emotional regions for its expression (acquisition and recall), and on spatial-contextual-mnemonic network for its persistence and (maybe) relapse.

### Connectivity on exposure to saline-associated cues is highly correlated with morphine-CPP test scoring

We next tested whether connectivity during contextual cue presentation would correlate with performance in the CPP test. Using the same strategy, we plotted the z-scored connectivity upon saline or morphine-associated cue presentation against individual CPP deltas. Of the same 45 pairs of regions in the putative cortico-limbic network, 13 exhibited a significant correlation between connectivity on context presentation and CPP delta. This was mainly found in the connections between amygdala, lateral septum, and episodic memory regions on exposure to saline-associated cues (Figure 6-a). Significant positive correlations were found in the LSI-LHb, LSI-RSC, VTA-CA1, and Cg-RSC pairs. Negative correlations were found in several anterior-cortical amygdaloid nucleus (ACo) connections, namely BMA, CeA, DG, and RSC. Connectivity on exposure to morphine-associated cues was only correlated with CPP delta in the LSI-Cg pair. Graph theoretical representations of the border animals are consistent with these findings, revealing high inter-individual variation in connectivity only on presentation of saline-associated contexts, where amygdala regions appear increasingly central in low CPP scoring rats, and memory-related regions, including the lateral septum, appearing highly central for the highest scoring rat (Figure 6-b, left graphs).

**Figure 6.**
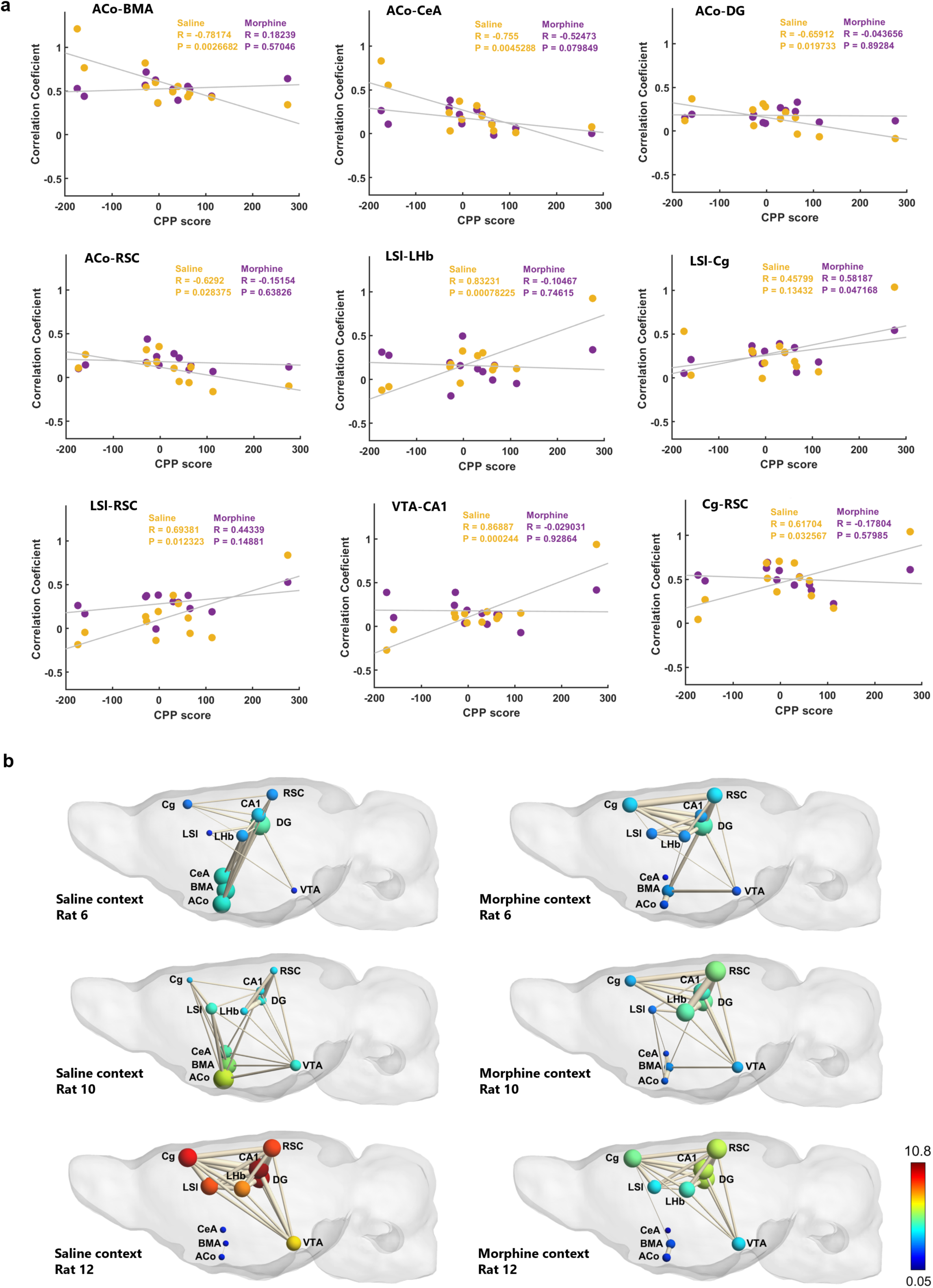
Connectivity during saline context exposure is the better predictor of morphine-CPP test scoring. a. Statistically significant correlation plots between connectivity during exposure to saline and morphine contexts in the indicated regions of the network and CPP delta. Values represent the CPP delta and the correlation coefficient for each individual animal in saline (yellow) and morphine (magenta) context conditions. R^2^ and p values are indicated in each panel. b. Graph theory representation of the putative cortico-limbic network during exposure to saline and morphine contexts for three border animals, two bad (rats 6 and 10) and one good learner (rat 12). Color scale represents node color, which was defined from a direct conversion of nodal degree into the jet colormap code in MATLAB (scale limits were set based on the least and most central nodes in our analysis).

CPP expression was uneven across animals, but the connectivity in response to morphine-associated cues in reward- and context-processing regions is similar, indicating that animals must have established a similar emotional association with the morphine-paired contexts. The connectivity in the same regions on exposure to saline-associated cues, on the other hand, is different among animals, indicating that different rats may have encoded different types of information across sessions of saline conditioning. In the saline-paired compartment, it is possible that a pre-anticipatory response to morphine administration and conditioning may have developed across sessions, stronger than the one developed during conditioning with morphine. It is also possible that a withdrawal/negative emotional state (due to absence of morphine) may have been encoded in the brain during saline conditioning. Our results align at the minimum with the first proposed explanation. As connectivity between ACo, a region implicated in odor-guided reward anticipation, and other contextual (RSC and DG) and emotional-related (BMA) regions was higher in animals that did not develop CPP, it is likely that they also encoded saline-associated cues as rewarding. Such overlap may have hampered their ability to fully dissociate the two contexts’ relevance and rewarding value. This is further supported by the lower connectivity in salience assignment and contextual encoding regions such as the VTA, septum, hippocampus, and cortex.

## Discussion

To investigate the brain-wide regional activity underlying drug addiction we have developed a morphine-induced CPP protocol whose contextual cues can be consistently presented inside an MRI apparatus. Our fMRI data led to the identification of several brain structures classically attributed to functional neural circuits responsible for specific cognitive functions. We found widespread BOLD signal changes in response to morphine- and saline-paired context cue presentation, namely regional circuitry involved in sensory processing (e.g. thalamus, somatosensory, piriform, visual, and auditory cortices), action planning and initiation (dorsolateral striatum and motor cortex (Balleine & O’Doherty, 2010)), and spatial cognitive and emotional processing (e.g. cortical: anterior cingulate, insular, retrosplenial, parietal and temporal associative cortices; subcortical: nucleus accumbens, lateral septum, hypothalamus, amygdala, nucleus reuniens, ventral tegmental area, dorsal raphe, and hippocampus).

Exposing rats to the sensory cues present in the distinct contexts led to BOLD responses in distinct regions according to context, suggesting that differentiation of context perception was achieved. In the presence of morphine-paired cues, we found activation of regions involved in the integration of limbic and mnemonic information (all PFC subdivisions, HIPP, DS, PRh, MEC and DLEnt (Laubach et al., 2018; Witter et al., 2017; Xu et al., 2016)), valence monitoring (BNST (Avery et al., 2016)), and arousal (AHA, PHA, p1RF). This agrees with immediate early gene expression data (Koya et al., 2006), showing PFC responses to cues associated to another opioid drug, heroin. Strong and extensive hippocampal responses confirm its established role in encoding, storing, and retrieving M-CPP memory (Portugal et al., 2014; Rivera et al., 2015), and indicate that the hippocampal formation is a potential neural substrate for the representation and storage of morphine-related memories. To our knowledge no previous studies have reported activation of the other regions in response to morphine-related contexts. Conversely, in the presence of saline-paired cues, we saw responses in regions involved in contextual (Calandreau et al., 2007) and emotional memory (Rogers & See, 2007): the medial septum (MS), ventral hippocampus (vHIPP), and periaqueductal gray (PAG) involved in avoidance of negative emotional states related to opioid withdrawal (Vázquez-León et al., 2021), with MS mediating aversiveness (via projections to the lateral habenula (LHb)) and locomotion (via projections to the preoptic area (POA)), for effective avoidance of uncomfortable environments (Zhang et al., 2018).

Here we also found extensive activation of the primary motor cortex, suggesting that saline-paired contexts were salient enough to engage different memory circuits, possibly due to a strong association formed between such context and the absence of morphine administration, the ensuing negative affective state (withdrawal), and supporting motor actions, ultimately reflecting motivation to avoid such discomfort. Since activity was primarily found in the first 30 sec of stimulus onset, it seems that perception, context recall, and possibly discrimination, occur immediately upon cue presentation. Whether this reflects a habituation process to an already recognized set of familiar stimuli remains to be determined.

Using network and seed-based connectivity analysis we found that, after successive drug-context pairings, the lateral septum (LSI) had increased its resting state functional connectivity with the regions of a putative cortico-limbic network. Besides being the strongest non-hippocampal target of hippocampal CA3 outputs, LSI plays a central role in modulating motivated behavior by integrating excitatory cortical inputs and regulating subcortical regions via tonic inhibition (Besnard & Leroy, 2022), namely through GABA-mediated inhibition over VTA during encoding and recall of M-CPP (Jiang et al., 2018; Luo et al., 2011). Enhanced LSI resting state functional connectivity is consistent with a role in regulating the encoding of contextual cues associated to morphine reward. Whether these changes reflect compensatory mechanisms to counter a strong dopaminergic output from VTA, such as induced by morphine, or to facilitate VTA dopaminergic signaling in the hippocampus remains an open question. Overall, the resting state functional connections within our predefined cortico-limbic network did not strikingly change after animals underwent a regime of morphine administration and conditioning. However, the connection between the lateral habenula and the CA1 region of the hippocampus was strengthened. Though not directly connected anatomically, the LHb maintains coherent theta oscillatory activity with dorsal hippocampus, directly related to memory performance in a spatial memory task (Goutagny et al., 2013). The LHb may, thus, play a role in encoding and consolidating contextual memories in the hippocampus, with either the drug or the contextual encoding, increasing synchrony. Another possible explanation for this increase in resting state connectivity may be related to the encoding of saline-related contexts. The LHb is known to regulate negative emotional affect, namely learning undesired experiences, and is also a critical component of an anti-reward system characterized by anticipation of aversive outcomes (in contrast with dopaminergic neuron activity) and an decrease in dopaminergic neuron firing in the VTA in response to failure to receive an expected reward (Hu et al., 2020; Koob & Volkow, 2016; Mirrione et al., 2014). It is possible that, as a result of consistent alternations between drug and saline conditioning, LHb activity is highly dysregulated. For example, after a single administration of cocaine, Jhou et al. (2013) found biphasic LHb responses, with an initial activity decrease followed by increase, consistent with an initial rewarding, and subsequent aversive, effect. Notably, this increase in activity persisted for 7 days following the last injection in a self-administration protocol. Although in our study the LHb did not significantly increase centrality in the putative cortico-limbic network (*p* = 0.0629), we cannot discard the possibility that, at the time of fMRI acquisition, animals were already entering the first stages of withdrawal, especially 4 hrs after having visited the M-CPP apparatus without receiving morphine. Altogether, the reinforced CA1-LHb connections and the largely consistent increase in lateral septum intra-network connectivity coincide with the processing, and likely consolidation, of emotionally charged episodic memories.

When morphine and saline-associated contexts are presented to the animals, a circuit formed by the amygdalás cortical and subcortical divisions, the habenula and septum presents itself as a central hub for contextual discrimination. Connectivity changes between the anterior cortical amygdaloid nucleus (ACo), a region seemingly involved in odor-guided reward anticipation (Shiotani et al., 2020), and the LSI, via opposed signal correlated activity in morphine- and saline-paired contexts, are consistent with the lateral septum playing the role of a reward effect modulator (or lever), during the encoding of relevant contextual cues. The central amygdala (CeA) and LHb also exhibit such opposed correlation dichotomy between morphine- and saline-paired contexts. We found anti-correlation during morphine context exposure, and vice-versa during saline context exposure. Ethanol addiction, binge-like intake of ethanol in mice, reveals enhanced activity of inhibitory projections from CeA to LHb (Companion et al., 2022). While we cannot ascertain the direction of this functional connection, and it is unclear whether negative correlations indeed reflect inhibitory processes at the cellular or circuit level (Becerra et al., 2006; Goelman et al., 2014), we consider plausible that the anti-correlation we found reflects the activity of such inhibitory synapse, resulting in silencing of antireward activity in LHb.

The level of functional connectivity between LHb and the three subdivisions of the amygdala, assessed during pre-CPP resting state fMRI, predicts individual M-CPP behavior. In rats with low CPP-delta we observed negative or near-zero correlation coefficients between these regions, while higher-scoring rats show positive coefficients. We did not find any correlation between CPP-delta scores and the connectivity within memory regions. Such correlations were specific to regions implicated in regulating negative affect, leading us to speculate that they reflect distinct circuit organization or inhibitory/excitatory connection imbalances among these regions across different animals. Furthermore, we find that connectivity levels within our limbic network were highly predictive of M-CPP scores during exposure to saline-associated cues, perhaps reflecting a parallel between the strength of CPP memory and the intensity of withdrawal-related neural circuit activity. The fact that, on exposure to morphine-paired cues, the levels of connectivity across regions involved in memory, reward, and affect, is not predictive of observed M-CPP deltas implies that neural functional connectivity is similar during morphine conditioning, and the differentiator factor might consist in either neural plasticity mechanisms at work during saline conditioning (S-CPP), as they preceded morphine conditioning in the same day, or the processing of short-intermittent withdrawal states throughout conditioning sessions. Further exploration is needed to understand whether low-scoring rats were protected from developing addiction due to being unable to encode or segregate different types of information in dedicated neural assemblies, or exhibited an intensified negative response to withdrawal that overshadowed their memory of morphine reward in the same context, or their motivation to approach the previously morphine-paired context. Alternatively, they may have established an aversive association with morphine administration and conditioning, attributing saline conditioning (absence of morphine) a rewarding value.

We have found a common circuitry supporting the neural mechanisms responsible for memorizing an association between distinct contexts, and morphine or the absence thereof. Such a common circuit includes regions involved in affect, reward, contextual perception and memory, with subtle, intriguing, functional specificities ultimately underlying the storage of distinct, individual, CPP memory engrams across the different animals. Animals that are more prone to strong emotional responses (as measured by the baseline resting state amygdala and habenula functional connectivity), exhibit a kind of neural circuit priming effect, even in the presence of saline contexts, and thus develop stronger connectivity patterns in anticipation of stronger morphine addiction behavior.

## Author contributions

JGR has designed the experiments, written the draft, performed all the experiments and data analysis, except the fMRI acquisition which was performed by JS. JM generated the MathLab codes for resting state and task-based fMRI analysis. SDG developed the algorithm for activity fMRI analysis. TS, JEC and MR helped in the CPP design implementation and analysis. MR, MCB, JEC, and LVL coordinated the project. All authors have revised the manuscript, discussed the experimental findings, and approved the manuscript.

## Acknowledgments

JGR is an FCT/PhD Fellow (IMM LisbonBioMed PhD program PD/BD/135518/2018); LVL is an FCT-CEEC Coordinator Investigator and is supported by FCT (2022.03699.CEECIND). TS is an FCT-CEEC Principal Investigator also supported by FCT (2022.03699.CEECIND). Funding from CIBIT was secured by FCT/UIDB&P/4950. We would also like to acknowledge the Rodent Facilities of Instituto de Medicina Molecular João Lobo Antunes and ICNAS for their technical support. We are also indebted to Marcelo Dias and Tiago Gil-Oliveira (ICVS, Braga), Patrícia Figueiredo (IST, Portugal), Disha Shah (VIB Leuven), Ingrid Bethus and Sebastian P. Fernandez (IPMC, Nice), and Gwenaelle Catheline (Univ. Bordeaux) for the availability and fruitful discussions/suggestions regarding data analysis. The graphical abstract was created with BioRender.com.

## Data availability statement

All the software used for data analysis and the respective information is provided in each respective section. The data that support the findings of this study are available from the corresponding authors upon reasonable request.

---

**Supplementary Figure 1.**
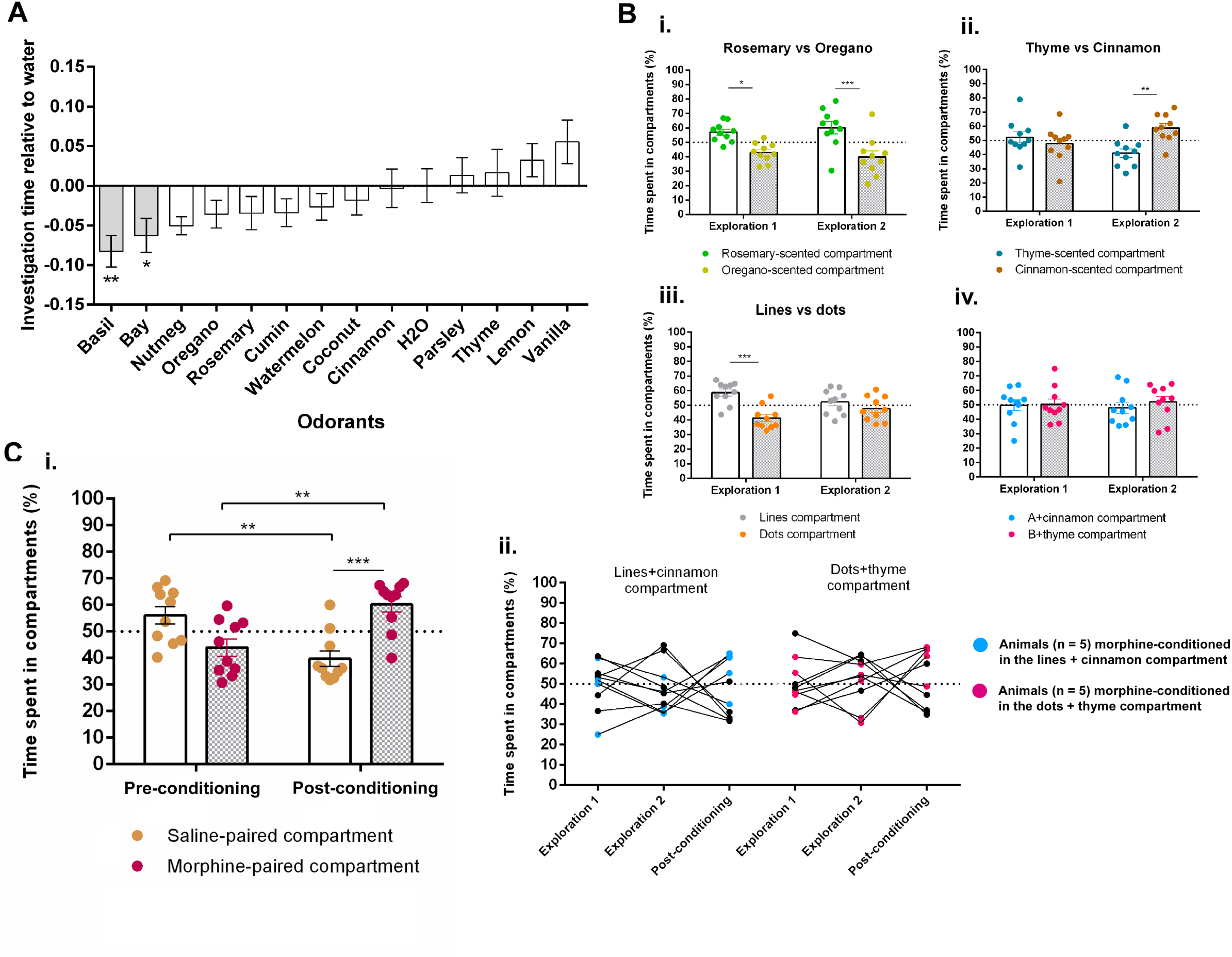
Validation of the contextual cues for fMRI adaptation of the CPP task. a. Spontaneous preference to indicated odorants relative to water (analysis of variance with post-hoc tests (Fisher LSD) performed on normalized investigation times). Values presented as mean ± SEM. *P < 0.05; **P < 0.01 b. Permanence times in compartments during a 15-min free exploration of the CPP apparatus under exposure to single odorants (i) and (ii); white light LEDs (array and single light point) (iii); and the combination of the best group of odorants and visual stimuli (iv). All values are presented as mean ± SEM. *P < 0.05; **P < 0.01; ***P < 0,001 (Two-way ANOVA followed by Bonferroni’s multiple comparisons test). c. i. Rats (n = 10) developed preference for the morphine-paired context, as revealed by the percentage of time spent in the morphine-paired compartment relative to time spent in the saline-paired compartment at post-conditioning, as well as between pre- and post-conditioning for both treatments. A two-way ANOVA followed by Bonferroni’s multiple comparisons test was employed to analyze differences between times spent in the two chambers. Values are presented as mean ± SEM. **P < 0.01, ***P < 0.0001. ii. Individual permanence times in the two CPP compartments in the pre- and post-conditioning sessions shows that M-CPP was expressed in most animals of the group (8 out of 10).

**Supplementary Figure 2.**
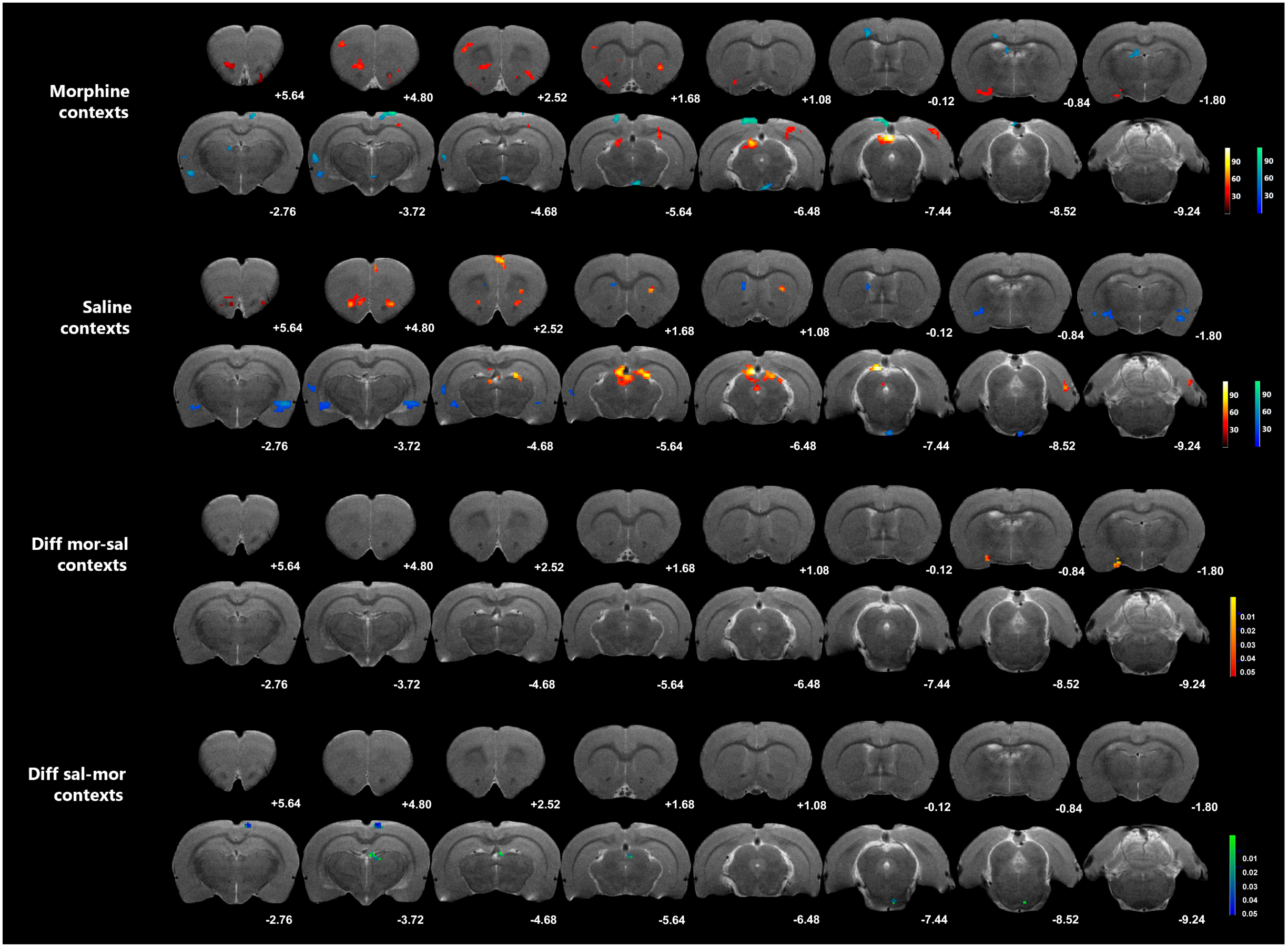
Responses to saline- and morphine-associated contexts by the end of stimulus presentation. Representative anatomical scans (T2-weighted anatomical MRI image of our template brain) display: i. and ii. Mean BOLD activity maps in the second-to-last 30-s window in response to the morphine- and saline-associated cues, respectively. Color scale represents mean regression coefficients for morphine and saline contrasts in comparison to baseline (masked under a significance level set at 0.05 and a cluster size threshold of 16 voxels); and iii. and iv. Magnitude maps show the clusters that survived the difference between activated clusters during presentation of saline and morphine cues, and vice-versa. Color scale represents P values of the surviving voxels. Coordinates under each brain section represent Paxinos & Watson’s rat brain atlas coordinates (in mm).

**Supplementary Figure 3.**
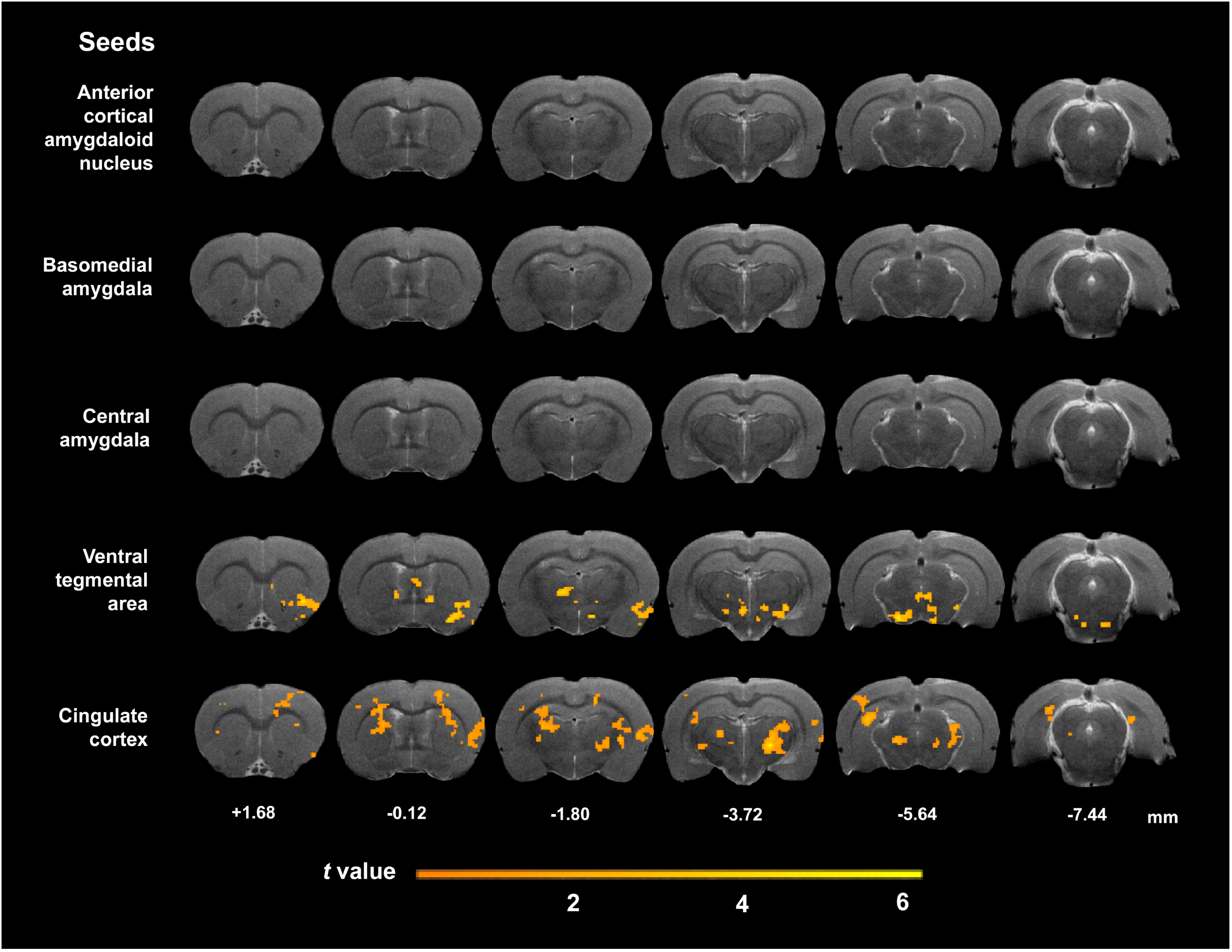
Resting state connectivity based on the seed-based analysis of the putative cortico-limbic network. Representative anatomical scans (T2-weighted anatomical MRI image of our template brain) display mean statistical t maps of resting state connectivity for the five supplementary regions within our putative cortico-limbic network, converted into seeds for seed-based analysis. Color scale represents t-value i.e., strength of functional connectivity of the seed region with all voxels in the brain. Abbreviations: ACo = anterior cortical amygdaloid nucleus, BMA = basomedial amygdala, CeA = central amygdala, VTA = ventral tegmental area, Cg = cingulate cortex. Coordinates under each brain section represent Paxinos & Watson’s rat brain atlas coordinates (in mm).

**Supplementary Figure 4.**
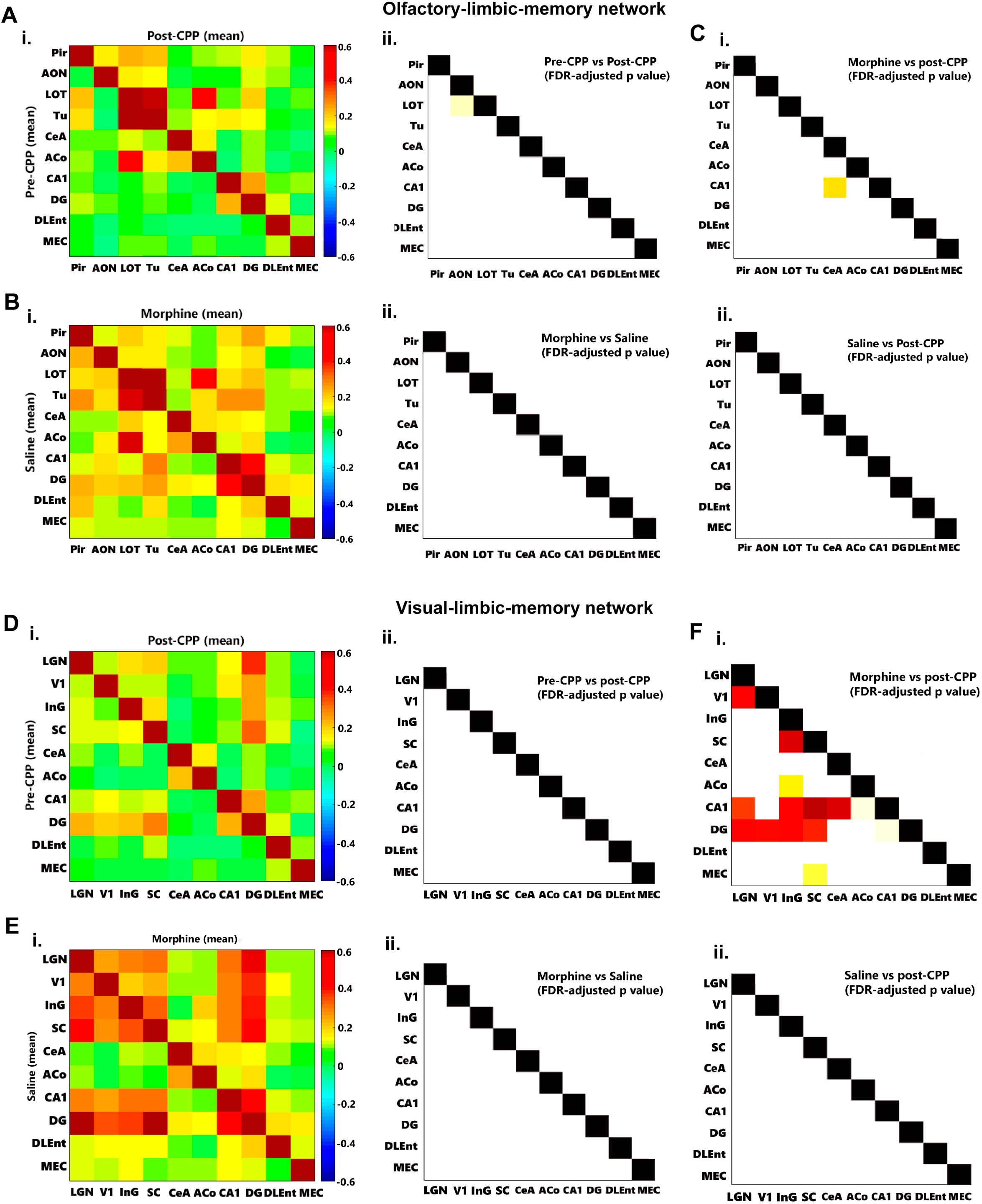
Functional connectivity in pre- and post-CPP, under saline and morphine context exposure for sensory and limbic regions. a. Functional connectivity (FC) between pre- and post-CPP in an olfactory-limbic-memory network. i. Upper panel: Mean FC-matrices of pre-CPP (lower half) and post-CPP (upper half) contexts. X and Y-axes represent brain regions. Color scale represents Pearson correlations i.e., strength of FC between each pair of brain regions. ii. FDR-adjusted P value matrix representing statistically significant connectivity differences between pre- versus post-CPP. Color scale represents P values (0-0.05). X- and Y-axes represent brain regions. b. Functional connectivity (FC) between saline- and morphine-paired contexts in an olfactory-limbic-memory network. i. Upper panel: Mean FC-matrices of saline (lower half) and morphine (upper half) contexts. X and Y-axes represent brain regions. Color scale represents Pearson correlations i.e., strength of FC between each pair of brain regions. ii. FDR-adjusted P value matrix representing statistically significant connectivity differences between saline versus morphine. Color scale represents p values (0-0.05). X- and Y-axes represent brain regions. c. FDR-adjusted P value matrices representing statistically significant connectivity differences between i. morphine versus post-CPP and ii. saline versus post-CPP. Color scale represents p values (0-0.05). X- and Y-axes represent brain regions. Abbreviations: Pir = piriform cortex, AON = anterior olfactory nucleus, LOT = nucleus of the lateral olfactory tract, Tu = olfactory tubercle, CeA = central amygdala, ACo = anterior cortical amygdaloid nucleus, CA1 = CA1 region of the hippocampus, DG = dentate gyrus, DLEnt = dorsolateral entorhinal cortex, MEC = medial entorhinal cortex. d. Functional connectivity (FC) between pre- and post-CPP in an visual-limbic-memory network. i. Upper panel: Mean FC-matrices of pre-CPP (lower half) and post-CPP (upper half) contexts. X and Y-axes represent brain regions. Color scale represents Pearson correlations i.e., strength of FC between each pair of brain regions. ii. FDR-adjusted P value matrix representing statistically significant connectivity differences between pre- versus post-CPP. Color scale represents P values (0-0.05). X- and Y-axes represent brain regions. e. Functional connectivity (FC) between saline- and morphine-paired contexts in an olfactory-limbic-memory network. i. Upper panel: Mean FC-matrices of saline (lower half) and morphine (upper half) contexts. X and Y-axes represent brain regions. Color scale represents Pearson correlations i.e., strength of FC between each pair of brain regions. ii. FDR-adjusted P value matrix representing statistically significant connectivity differences between saline versus morphine. Color scale represents p values (0-0.05). X- and Y-axes represent brain regions. f. FDR-adjusted P value matrices representing statistically significant connectivity differences between i. morphine versus post-CPP and ii. saline versus post-CPP. Color scale represents p values (0-0.05). X- and Y-axes represent brain regions. Abbreviations: LGN = lateral geniculate nucleus, V1 = primary visual cortex, InG = intermediate gray layer of the superior colliculus, SC = superior colliculus, CeA = central amygdala, ACo = anterior cortical amygdaloid nucleus, CA1 = CA1 region of the hippocampus, DG = dentate gyrus, DLEnt = dorsolateral entorhinal cortex, MEC = medial entorhinal cortex.

